# The fitness landscape of the African *Salmonella* Typhimurium ST313 strain D23580 reveals unique properties of the pBT1 plasmid

**DOI:** 10.1101/689075

**Authors:** Rocío Canals, Roy R. Chaudhuri, Rebecca E. Steiner, Siân V. Owen, Natalia Quinones-Olvera, Melita A. Gordon, Michael Ibba, Jay C. D. Hinton

## Abstract

We have used a transposon insertion sequencing (TIS) approach to establish the fitness landscape of the African *Salmonella enterica* serovar Typhimurium ST313 strain D23580, to complement our previous comparative genomic and functional transcriptomic studies. We used a genome-wide transposon library with insertions every 10 nucleotides to identify genes required for survival and growth *in vitro* and during infection of murine macrophages. The analysis revealed genomic regions important for fitness under two *in vitro* growth conditions. Overall, 724 coding genes were required for optimal growth in LB medium, and 851 coding genes were required for growth in SPI-2-inducing minimal medium. These findings were consistent with the essentiality analyses of other *S.* Typhimurium ST19 and S. Typhi strains. The global mutagenesis approach also identified 60 sRNAs and 413 intergenic regions required for growth in at least one *in vitro* growth condition. By infecting murine macrophages with the transposon library, we identified 68 genes that were required for intra-macrophage replication but did not impact fitness *in vitro*. None of these genes were unique to *S.* Typhimurium D23580, consistent with a high conservation of gene function between *S.* Typhimurium ST313 and ST19 and suggesting that novel virulence factors are not involved in the interaction of strain D23580 with murine macrophages. We discovered that transposon insertions rarely occurred in many pBT1 plasmid-encoded genes (36), compared with genes carried by the pSLT-BT virulence plasmid and other bacterial plasmids. The key essential protein encoded by pBT1 is a cysteinyl-tRNA synthetase, and our enzymological analysis revealed that the plasmid-encoded CysRS^pBT1^ had a lower ability to charge tRNA than the chromosomally-encoded CysRS^chr^ enzyme. The presence of aminoacyl-tRNA synthetases in plasmids from a range of gram-negative and gram-positive bacteria suggests that plasmid-encoded essential genes are more common than had been appreciated.

## Introduction

*Salmonella* spp. are important pathogens of humans and animals. In humans, salmonellosis is classified as either a typhoidal or non-typhoidal *Salmonella* (NTS) disease. Typhoidal salmonellosis involves systemic spread through the body that causes enteric fever, and is associated with the *S. enterica* serovars Typhi (*S.* Typhi) and Paratyphi (S. Paratyphi). In contrast, NTS disease normally involves a self-limiting gastroenteritis that is transmitted via food, involving approximately 94 million human cases and about 155,000 deaths [1]. The *S. enterica* serovar Typhimurium (*S*. Typhimurium) sequence type ST19 causes the majority of gastroenteritis in immuno-competent individuals worldwide via pathogenic mechanisms that induce mucosal inflammatory responses in the gut. *S.* Typhimurium can thrive in this inflamed gut whilst other key members of the gut microbiota cannot survive [2, 3]. The remarkable ability of this pathovariant to enter, survive, and proliferate in mammalian macrophages and epithelial cells in a “*Salmonella*-containing vacuole” (SCV) is responsible for systemic disease in both animals and humans [4].

The HIV epidemic in sub-Saharan Africa has been implicated in the evolution of new clades of NTS strains that cause bacteraemia in humans. Specifically, the HIV virus impairs the immunity of adults, a phenomenon that occurred concurrently with the development of NTS strains able to cause a systemic disease, invasive non-typhoidal salmonellosis (iNTS) [5–9]. In children, malaria and malnutrition are also risk factors for iNTS [10]. The *S.* Typhimurium and *S.* Enteritidis isolates responsible for invasive NTS isolates have a multi-drug-resistant phenotype, necessitating the replacement of conventional therapies with alternative antibiotics [6,11,12].

In sub-Saharan Africa, *S.* Typhimurium strains belonging to sequence type ST313 have been associated with the majority of systemic disease, causing hundreds of thousands of deaths in 2010 [13]. The genome sequence of one representative of ST313, D23580, was published in 2009 [14], and was recently updated [15].

To date, genome-wide functional genomic studies have focused on the fitness of *S.* Typhimurium and *S*. Typhi in several *in vitro* growth conditions, and within eukaryotic cells and animal infection models [16]. The recent development of transposon-insertion sequencing (TIS) technology combines global mutagenesis and high-throughput sequencing to functionally characterize bacterial genes. Transposon insertion libraries are constructed in a strain of interest, in which nonessential genes for a particular growth condition (input library) are dispensable and contain insertions. This library of random transposon insertion mutants can be used to identify genes “required” for fitness under that particular environmental condition (output library). The relative proportion of each mutant in the input and the output libraries is determined by high-throughput sequencing, a strategy that enables the fitness contribution of each gene to be quantified in environmental conditions of interest [17, 18]. The first study that used this technology in *Salmonella* described a new TIS strategy: Transposon-Directed Insertion Site Sequencing (TraDIS) [19]. Subsequently, various TIS-based strategies have been used for functional genomic analysis of *Salmonella* serovars Typhimurium and Typhi [20–22].

Here we report the TIS-based identification of the genes of *S*. Typhimurium ST313 D23580 responsible for growth and survival inside murine macrophages, and the genetic requirements of this strain to grow and survive in laboratory conditions (summarized in Figure 1).

**Figure 1.**
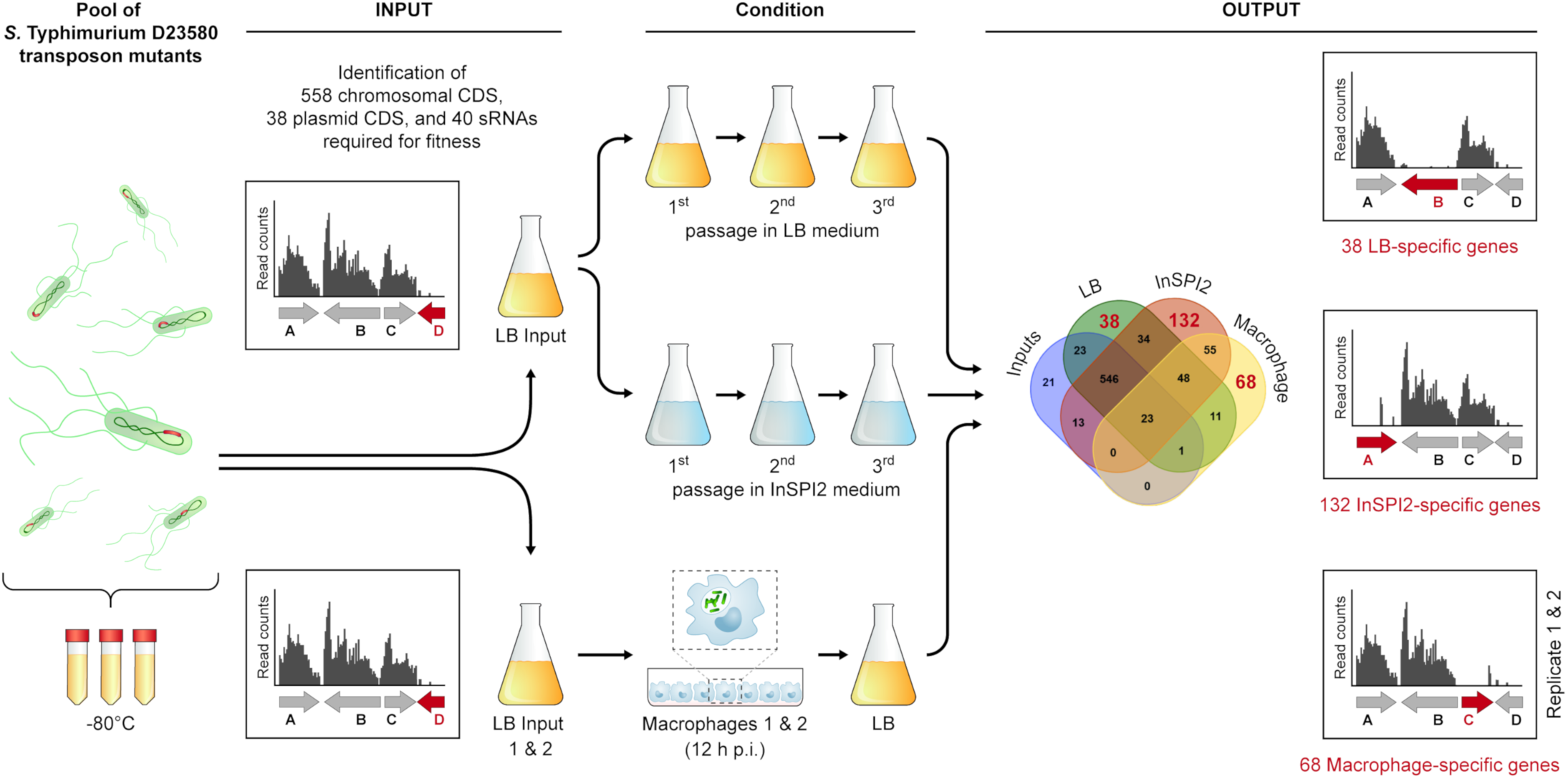
Transposon-insertion sequencing (TIS) in S. Typhimurium ST313 D23580. Schematic representation of the S. Typhimurium D23580 transposon library and growth conditions used in this study.

## Results and Discussion

### Transposon insertion profile of a *S*. Typhimurium D23580 Tn5 library

A transposon library was constructed in the *S*. Typhimurium ST313 lineage 2 representative strain D23580. The pool of transposon mutants was grown in LB (input) and successively passaged three times in two different laboratory growth media: a rich medium, LB (output); and an acidic phosphate-limiting minimal medium that induces *Salmonella* pathogenicity island (SPI) 2 expression, designated InSPI2 (Figure 1). Genomic DNA from the input and output samples was purified and prepared for Illumina sequencing of the DNA adjacent to the transposons (Materials and Methods). Table S1 shows the number of sequence reads obtained, the sequence reads that contained the transposon tag sequence, and the sequence reads that were uniquely mapped to the *S*. Typhimurium D23580 genome.

Sequence analysis of the input sample identified 797,000 unique transposon insertion sites in *S*. Typhimurium D23580, equating to an average of one transposon integration every six nucleotides. All data are available for visualization in a Dalliance genome browser [23] which shows the transposon insertion profile of the chromosome and the four plasmids (pSLT-BT, pBT1, pBT2, pBT3) in *S*. Typhimurium D23580: https://hactar.shef.ac.uk/D23580. The number of reads, transposon insertion sites, insertion index, and “requirement” call per gene are summarized in Table S2. The insertion index was calculated as described in Materials and Methods and allowed the genetic requirements of *S*. Typhimurium D23580 for growth to be determined after a single passage in LB. The genes designated “required” included essential genes and genes that contributed to fitness in this particular environmental condition. Some genes were called “ambiguous” when they could not be robustly assigned as either required or dispensable by the analysis (Materials and Methods). A total of 596 genes were required in the *S*. Typhimurium D23580 genome: 558 were located in the chromosome, two in the pSLT-BT plasmid, and 36 in the pBT1 plasmid (Figures 2A and B).

**Figure 2.**
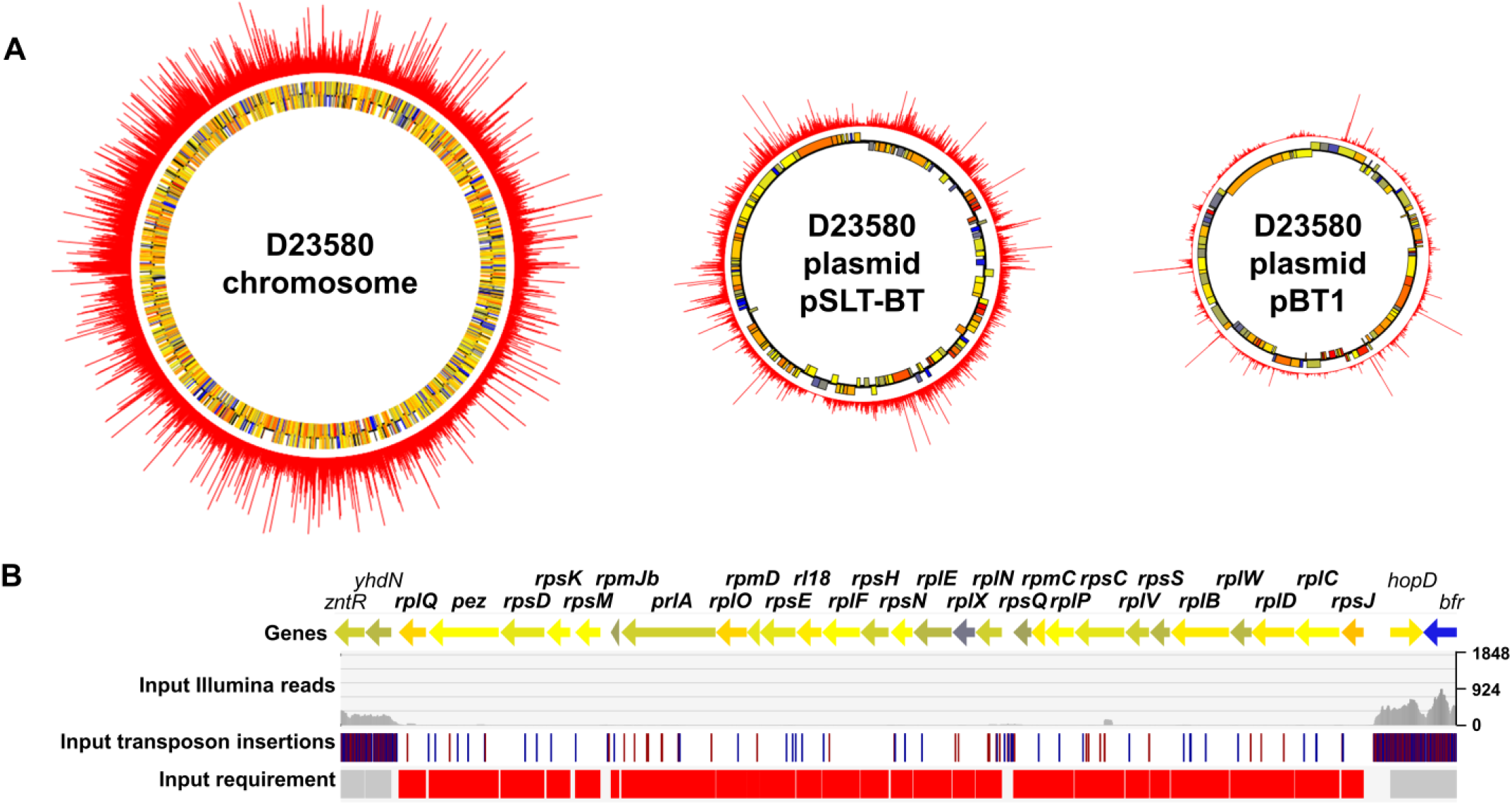
Transposon insertion profile in S. Typhimurium D23580. (A) Transposon insertion indexes are represented in the outer ring of the chromosome and the pSLT-BT and pBT1 plasmids of *S*. Typhimurium D23580. The two inner rings represent annotated genes coloured according to their GC content (blue = low, yellow = intermediate, red = high). (B) Chromosomal region showing a cluster of genes, between *zntR-yhdN* and *hopD-bfr*, that are required for growth in D23580.

To establish common themes amongst closely-related bacteria, the *S*. Typhimurium D23580 chromosomal genes that were required for growth were compared with the genetic requirements of other strains of *S*. Typhimurium (14028, a derivative of SL1344 called SL3261, and LT2) and *S*. Typhi (Ty2, and a derivative of Ty2 named WT174) (Figure 3A, Table S3) [19,20,24–26]. After the comparison with these five other *Salmonella* isolates, we found that a total of 101 genes were only required in D23580, including one D23580-specific gene encoding the CI^BTP5^ repressor of the BTP5 prophage (*STMMW_32121*) [27]. To add context, a Clusters of Orthologous Groups (COG) analysis identified 32 genes that were predominantly assigned to two functional categories of transcription (nine genes) and amino acid transport and metabolism (six genes). Additionally, at least 21 of the 101 genes were associated with virulence: 16 were located in *Salmonella* pathogenicity island (SPI) regions, and five encoded associated effectors that were located elsewhere in the genome.

**Figure 3.**
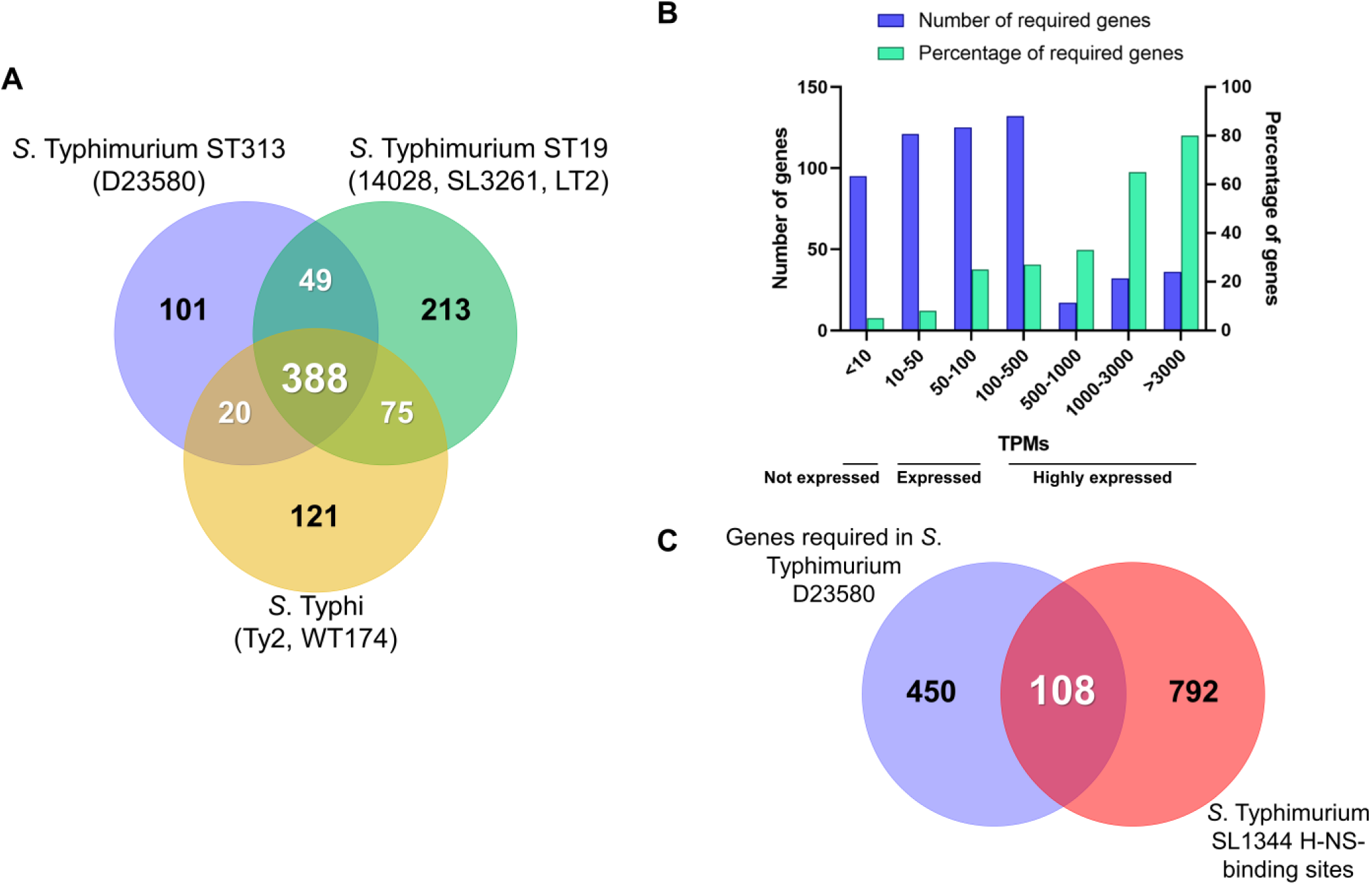
*S*. Typhimurium D23580 chromosomal genes required for growth *in vitro*. (A) *S*. Typhimurium D23580 required chromosomal genes for growth that have never been previously identified as required for growth in *S*. Typhimurium (*S*. Typhimurium 14028 [20], SL3261 [24], 14028 [74], LT2 [26]), or *S*. Typhi (Ty2 [20], WT174 [19]). For the previously published studies, only genes that shared an ortholog in D23580 were used for the final analysis. (B) Transcriptional levels of expression in LB mid-exponential phase (MEP) of the D23580 required genes for growth (data extracted from Canals and colleagues [15]). The X axis represents groups of absolute TPM values. The Y axis represents the number of required genes in D23580. The second Y axis represents the percentage of required genes with a TPM value within the range showed in the X axis out of the total number of genes showing a TPM value within the same range. (C) Required genes in D23580 with H-NS-binding sites reported in *S*. Typhimurium SL1344 [36]. Only the SL1344 genes that shared an ortholog with D23580 were included for the analysis.

To identify chromosomal genes that are required for growth in *S*. Typhimurium D23580 and other *S*. Typhimurium pathovariants, a comparison with the individual strains was performed (Figure S1A, Table S4) [20, 24]. A total of 250 genes were required in all *S*. Typhimurium and *S*. Typhi strains [19,20,24]. While searching for serovar-specific required genes, we found six genes that were only required by D23580 and the two *S*. Typhimurium strains 14028 and SL3261, but not by *S.* Typhi, namely: *ssaT*, a SPI-2 gene; *STMMW_16291*, encoding a putative amino acid transporter; *hnr*, encoding a regulator; *pth*, encoding a peptidyl-tRNA hydrolase; *STMMW_18451*, with unknown function; and *ddhB* (*rfbG*), an O-antigen gene involved in the biosynthesis of CDP-abequose. Intriguingly, *hns* was only required in *S*. Typhimurium D23580 and 14028 but not in SL3261. In contrast, no genes that were only required by *S*. Typhimurium D23580 and the two *S*. Typhi strains Ty2 and WT174 were found. We conclude that there is a high level of conservation of genes that contribute to fitness during *in vitro* growth of *S*. Typhimurium and *S*. Typhi. Because D23580 is much more closely related to other strains of *S*. Typhimurium than to *S*. Typhi, it is not surprising to see that there is a greater overlap of required genes between the Typhimurium strains than with the Typhi isolates.

Seven genes previously reported to be required in *S*. Typhimurium were not included in the D23580 gene requirements (Figure S1B, Table S4), prompting a more detailed investigation. Analysis of two other input samples described later in this work showed that two of these genes were consistently identified as dispensable in D23580: *cysS*, encoding a cysteinyl-tRNA synthetase; and *folA*, involved in biosynthesis of tetrahydrofolate. There were 20 genes that had been previously reported in TIS studies to be required in *S*. Typhi but not in D23580 (Figure S1C, Table S4) [19, 20]. Seven out of the 20 genes were consistently found to be dispensable in the two other D23580 input samples analyzed later in this work, and one of them was *cysS*. The cysteinyl-tRNA synthetase is essential for bacterial growth [28]. The dispensability of *cysS* (*cysS^chr^*) in D23580 reflects the fact that the pBT1 plasmid of *S*. Typhimurium D23580 carries the paralogous gene *cysS^pBT1^*, and we published the transposon insertion profiles of these two genes previously [15]. In summary, only the plasmid copy *cysS^pBT1^* was required for growth in LB in D23580, whereas *cysS^chr^* was dispensable.

### Certain dispensable genes were designated required in the essentiality analysis

Several genes required for *in vitro* growth of D23580 have previously been reported to be dispensable for growth in other *Salmonella* strains in laboratory conditions, including 12 genes located in the SPI-2 pathogenicity island and six genes in the O-antigen biosynthetic cluster. Here, the low number or absence of transposon insertions could reflect a limitation of the TIS technique. Although a previous study using a similar strategy for Tn5 transposon library construction did not find bias in the insertion sites [20], a preference of Tn5 for G/C pairs in the target sequence has been reported [29]. Motifs for preferential Tn5 integration have been investigated previously [20, 30].

Searching for transposon insertion bias in previously published work, we found that the transposition of the Mu element was lower in highly transcribed regions of the chromosome [31, 32]. To correlate the level of transcription with the number of Tn5 transposon insertions, we assessed the absolute expression values of the required genes using our published *S*. Typhimurium D23580 RNA-seq data, grown in LB to mid-exponential phase (MEP) (Figure 3B, Table S3) [15]. In this dataset, the level of expression of each gene is expressed as Transcripts Per Million (TPM). We found that 80% of the most highly expressed genes (TPM >3000) were required for growth. Of these, 86% were involved in translation, ribosomal structure and biogenesis. We conclude that fewer Tn5 transposon insertions occurred in highly expressed genes and most of these genes are involved in functions that are essential for bacterial growth.

The histone-like nucleoid structuring (H-NS) protein, encoded by the *hns* gene, preferentially binds to A/T-rich regions in bacterial genomes [33, 34]. It has been proposed that H-NS-bound DNA could be protected from being a target of transposition and so receive fewer transposon integrations [35]. To investigate whether H-NS-binding explained the low number of transposons found in *Salmonella* genes that are dispensable for growth in laboratory conditions, the reported H-NS-binding sites of *S*. Typhimurium SL1344 [36] were cross-referenced with the list of required genes in D23580 (Figure 3C, Table S3). A total of 108 genes designated as required in D23580 also contained an H-NS binding site in SL1344 (Table S3), including 36 genes located in SPI regions and associated effectors. We conclude that only a minority of *S*. Typhimurium D23580 genes are likely to be protected from transposition by H-NS, as discussed below.

Transposon insertions were seen more rarely in SPI pathogenicity island-related *S*. Typhimurium D23580 genes than other parts of the chromosome. Specifically, the *hilC* (SPI-1) and *ssrA* (SPI-2) genes were designated as required in our study. To investigate whether the deletion of these genes had a fitness cost for D23580, we compared the growth of the individual D23580 deletion mutants with the D23580 wild-type (WT) strain in LB (Figure S2A, Tables S4 and S5). Similar mutants that retained the kanamycin (Km) resistance cassette were also examined, in case the strong promoter of the *aph* gene played a role. The deletion of *ssrA* included the removal of *ssrB*, the two genes encode the two-component regulatory system of SPI-2. Two mutants that lacked genes involved in the biosynthesis of the lipopolysaccharide (LPS), in *waaL* (lack of O-antigen) and *waaG* (absence of O-antigen and part of the LPS core), were also investigated as examples of genes containing H-NS-binding sites and allowing a high proportion of transposon insertions.

No significant differences were observed in the growth rate of any of the SPI-2 and the SPI-1-defective mutants compared to the WT strain. We observed that the SPI-1 mutant grew to a slightly greater culture density than the WT strain (OD_600_ for the WT was 4.55, and was 4.86 for D23580 Δ*hilC*::*frt*). This small fitness cost of expressing *hilC* and other SPI-1 genes has already been reported in *S*. Typhimurium, explaining why SPI-1 mutants outcompete the WT strain [37, 38]. Both LPS mutants grew slower in LB compared to WT, consistent with previous findings concerning the deletion of the *S*. Typhimurium *waaL* gene [39]. These results suggest that the binding of the H-NS protein to the SPI-1 and SPI-2 regions could explain the low number of transposons in these regions, as no fitness cost for growth in LB of the respective mutants was detected. The high number of transposon insertions found in the LPS genes, which were reported to contain H-NS-binding sites in *S*. Typhimurium SL1344, and the fact that mutations in these genes had an effect on fitness, indicated that a low number or absence of transposon insertions do not always correlate with the presence of H-NS.

### Genetic requirements for *S*. Typhimurium D23580 growing in rich and SPI-2-inducing media

To build on our previous identification of genes required for survival after a single passage in LB, we studied *in vitro* fitness during growth in nutrient rich and minimal media by comparing the pools of transposon mutants recovered after further passages of the D23580 transposon library in LB (designated as “output”), and the acidic phosphate-limiting minimal medium (PCN, phosphate carbon nitrogen) that induces SPI-2 expression (InSPI2) in *S. enterica.* These data were used to assign an insertion index to each gene (Table S2).

After three passages in LB, 724 genes were required, which included essential genes and genes that contributed to fitness for *in vitro* growth: 683 genes in the chromosome, 2 genes in the pSLT-BT plasmid, and 39 genes in the pBT1 plasmid. A total of 851 genes were required for optimal growth after three passages in InSPI2: 816 genes in the chromosome, 2 genes in the pSLT-BT plasmid, and 33 genes in the pBT1 plasmid. Genes that had previously been found to be indispensable for growth in the input sample in this study were removed from the lists of required genes (Figure 4A, Table S3).

**Figure 4.**
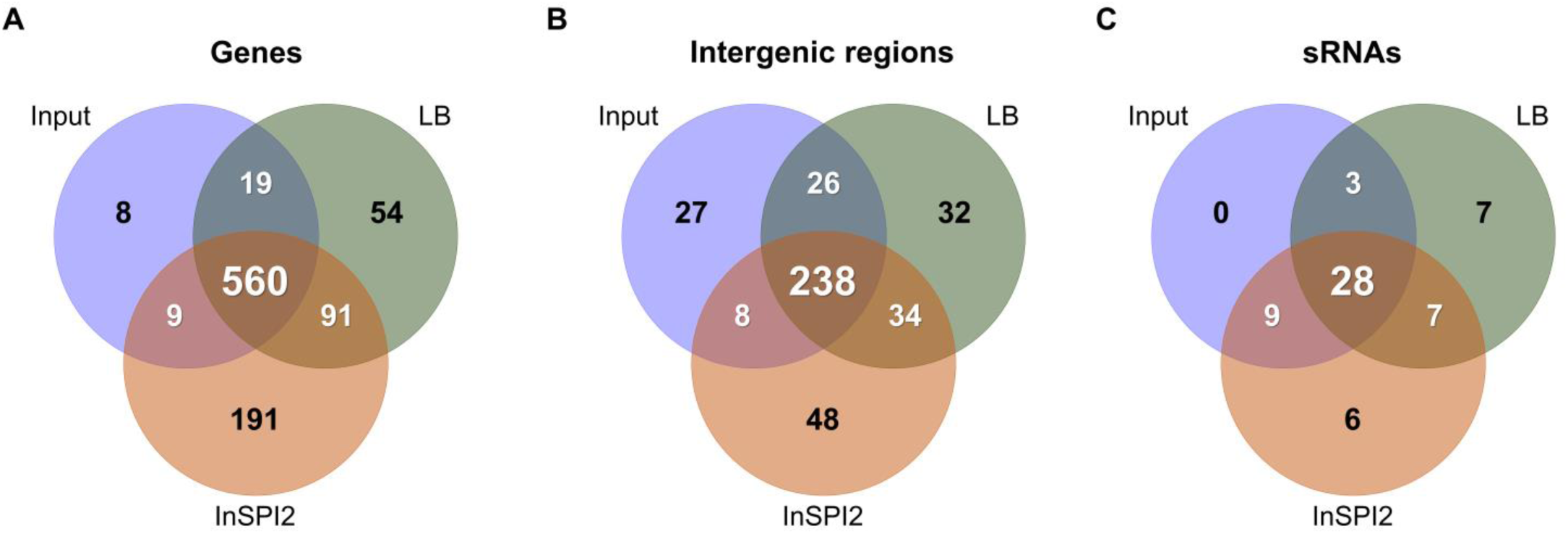
Identification of S. Typhimurium D23580 required for *in vitro* growth. The figures represent (A) coding genes; (B) intergenic regions; and (C) sRNAs.

There were 54 *S*. Typhimurium D23580 genes that were required for growth after three passages in LB, but not in InSPI2, including the pBT1-encoded *pBT1-0401* and *pBT1-0781*. The gene list included two genes reported to be required by *S*. Typhimurium 14028 after three passages in LB [21], namely *sdhA*, encoding a succinate dehydrogenase flavoprotein subunit; and *sapG* (*trkA*), encoding a potassium transporter. The identification of these particular *S*. Typhimurium genes in multiple TIS studies highlights the importance of *sdhA* and *sapG* for fitness in this environment. The genes *sdhCD*, involved in the conversion of succinate to fumarate (with *sdhAB*), and *fumA*, involved in the conversion of fumarate to malate (with *fumBC*), were also required after three passages in LB in our *S*. Typhimurium D23580 study. Overall, most of the required genes were either involved in energy production and conversion (17%), carbohydrate transport and metabolism (9%) or inorganic iron transport and metabolism (9%).

A total of 191 genes were needed for optimal growth of *S*. Typhimurium D23580 after three passages in the InSPI2 minimal medium. These genes included: *dksA*, that is required for growth on minimal medium [40]; *hfq*, encoding an RNA chaperone that facilitates base-pairing of ∼100 small RNAs (sRNAs) to the their target mRNAs [41, 42]; and genes involved in thiamine, coenzyme A, biotin, and LPS biosynthesis. The majority of the genes were included in the following functional categories: amino acid transport and metabolism (27%), cell envelope biogenesis and outer membrane (7%), inorganic iron transport and metabolism (7%), and nucleotide metabolism (7%).

There were 91 genes designated required in both LB and InSPI2 (Figure 4A, Table S3). Among them, we found two D23580-specific genes: *pBT1-0081*, in the pBT1 plasmid; and *cI^BTP1^*, encoding the BTP1 prophage repressor [27]. Most of the genes encoded products involved in energy production and conversion (22%), oxidative phosphorylation (12%), and translation, ribosomal structure and biogenesis (10%). These genes reflect the biological needs of *S*. Typhimurium D23580 for growth in laboratory conditions.

### The intergenic regions and sRNAs that increase fitness of *S*. Typhimurium D23580

The high level of saturation of the transposon library allowed the transposon insertion profiles of sRNAs and intergenic regions of ≥100 bp in the chromosome, and the pSLT-BT and pBT1 plasmids, to be investigated to identify the role of these regions in fitness [14, 15]. The pBT2 (∼2.5 kb) and pBT3 (∼2 kb) plasmids were considered as intergenic regions due to the absence of annotation [15]. An insertion index was calculated for each intergenic region and sRNA in the input and LB and InSPI2 output samples as previously described (Table S6).

A total of 286 intergenic chromosomal regions, 13 intergenic plasmid regions (all in pBT1), and 40 sRNAs were required in the input sample, being essential or contributing to fitness. Thirty-two intergenic regions were important for fitness after three passages in LB, while 48 were only required for growth after three passages in InSPI2 (Figure 4B, Table S3). Additionally, 34 intergenic regions increased fitness in both LB and InSPI2. Most of the adjacent genes of the intergenic regions that were important for fitness were involved in translation, ribosomal structure and biogenesis (11%), and energy production and conversion (9%). The coding genes required for optimal growth after three passages in LB and InSPI2 also belonged to these two functional categories (Table S2). The fitness defects of mutants carrying transposon insertions in intergenic regions might be due to disruption of the promoter region of one of the flanking genes, or could reflect mutation of unannotated coding regions.

Seven sRNAs were only required for growth after three passages in LB, but not in InSPI2 minimal media (Figure 4C, Table S3). In contrast, six sRNAs were identified as being important for fitness after three passages in InSPI2. In total, seven sRNAs that enhanced fitness in both LB and InSPI2 were identified, namely tp2, STnc2010, RyjB, STnc2030, SdsR, STnc3080, and SraG. Among them, SdsR is widely conserved in enterobacteria [43], targeting important global regulators with biological relevance in stationary phase and stress conditions [44] and shown to be required for fitness of *S.* Typhimurium in stationary phase [45]. The fact that inactivation of SdsR has already been described to have a fitness cost in stationary phase helps to validate our TIS approach. For the future, the regulatory targets of the other six sRNAs that are required for growth in both nutrient and minimal media should be determined.

### Intra-macrophage infection with the transposon library reveals the absence of novel virulence factors in *S*. Typhimurium D23580

To build upon our understanding of the *S.* Typhimurium D23580 genes that were required for fitness during growth in nutrient or minimal media, we used the transposon library to investigate the process of intracellular infection of murine RAW264.7 macrophages. To accurately identify genes required for growth with a transposon library, it is important to sequence the input sample as well as the output sample used for each experiment. As confirmed earlier, every time a transposon library is passaged in LB medium, mutants that exhibit reduced fitness will be lost from the library. Therefore, we sequenced two biological replicates of the library that had been grown in LB prior to infection of macrophages (Input 1 and Input 2 samples). Proliferation of the transposon library was assessed after a single passage through murine macrophages, as additional passages caused the selection of LPS mutants (Text S1, Figure S3). Accordingly, at 12 h post-infection (p.i.), intracellular bacteria from the two biological replicates were recovered (Macrophage 1 and Macrophage 2 samples) (Figure 1, Text S1). Genomic DNA from the two input and the two output samples was purified and Illumina sequenced. Table S1 contains the number of demultiplexed reads, the number of reads with a transposon tag, and the number of uniquely mapped reads. The two input samples contained 660,000 and 928,000 unique insertion sites. Combining these results with the analysis of the previous input dataset (797,000 unique insertion sites), a total of 511,000 unique insertion sites were common to all three datasets, an average of one transposon integration per ten nucleotides.

The majority of genes (87%) that were designated “required” in the two new inputs were consistent with the analysis of the previous input sample (Figure S4A, Table S4). Eleven of the genes considered ambiguous in the previous input sample were required in Input 1 and Input 2, including *cI^BTP1^*, the repressor of the D23580-specific prophage BTP1 [27]. Prophage repressors are commonly found to be required genes for growth in TIS studies because inactivation leads to prophage de-repression and phage-mediated cell lysis. Among the five complete prophage regions in D23580 (BTP1, Gifsy-2^D23580^, ST64B^D23580^, Gifsy-1^D23580^, and BTP5) [27], only the repressors of the two D23580-specific prophages, BTP1 and BTP5, were designated as required in our analyses.

The data were used to identify genes that contributed to fitness of *S.* Typhimurium D23580 during macrophage infection. Specifically, genes important for intracellular growth and survival in murine macrophages were identified by comparing the two macrophage output samples with the two input samples (Materials and Methods). Transposon insertions in 206 D23580 genes caused attenuation in the macrophage infection model (log2 fold-change <-1, *P*-value <0.05) (Table S7). Many of these genes correlated well with previous high-throughput studies of *S*. Typhimurium ST19 in different animal infection models (Figures S5A and S5B), including five well-characterized regulatory systems that control *Salmonella* virulence: the *phoPQ* two-component regulators; the *ssrAB* regulators of SPI-2 gene expression; *dam*, DNA adenine methylase; *hfq*; and *ompR*, an element of the two-component regulatory system *ompR-envZ*. Three D23580-specific genes, two in the pBT1 plasmid (*pBT1-0081* and *pBT1-0401*) and *cI^pBT1^*, had already been identified as important for fitness after three passages in LB in our study.

Inactivation of six D23580 genes increased fitness in the macrophage infection model (log2 fold-change >1, *P*-value <0.05): *nadD*, encoding a nicotinate-nucleotide adenylyltransferase; *STM1674* (*STMMW_16691*), encoding a transcriptional regulator; *barA*, encoding the sensor of the two-component regulatory system SirA/BarA that controls carbon metabolism via the CsrA/CsrB regulatory system; the pSLT-BT plasmid gene *repC;* and the LPS O-antigen biosynthetic genes *abe* (*rfbJ*) and *rmlA* (*rfbA*). Because the *abe* and *rmlA* mutants have short LPS which increases the invasiveness of *S.* Typhimurium without affecting intracellular replication [46], it is clear that our TIS strategy in macrophages not only identified mutants with increased fitness in terms of growth and survival inside macrophages but also selected for mutants that are more invasive in the infection model (Text S1).

### Macrophage-specific *S*. Typhimurium D23580 genes

To identify genes important for fitness inside macrophages but not for growth in laboratory media, the 206 D23580 genes that showed attenuation in macrophages when disrupted by a transposon insertion were cross-referenced with genes required for growth in the *in vitro* laboratory conditions tested in this study, LB and InSPI2 (Figure S4B, Table S4). We identified 182 “macrophage-associated genes” and, within this group, 68 “macrophage-specific genes” that had reduced fitness during macrophage infection and did not impact upon growth *in vitro*. The macrophage-specific genes included known *Salmonella* virulence genes: *phoPQ*, SPI-2 genes, *dam*, *hfq*, and *ompR*. Most of the 68 genes were involved in functions related to transcription (10%), amino acid transport and metabolism (9%), and translation, ribosomal structure and biogenesis (7%).

Analysis of our intra-macrophage transcriptome of *S.* Typhimurium ST19 showed that genes that encoded key virulence factors were macrophage-up-regulated by >3-fold [47]. Our recent D23580 RNA-seq results [15] led us to investigate the function of two genes in the macrophage infection model, *STM2475* and *STM1630*. Only the *STM2475* deletion mutant exhibited decreased intracellular replication, suggesting a putative role in virulence of D23580 (Text S1, Figures S6 and S7).

We previously showed that many *S.* Typhimurium virulence genes were both up-regulated within macrophages and required for animal infection [48]. We built on this concept by finding the “macrophage-specific” genes identified by transposon mutagenesis that were also up-regulated within macrophages, using transcriptomic data from Canals and colleagues [15] (Figure 5A). The results showed that the 23 genes that were required for intra-macrophage proliferation and were significantly “macrophage-up-regulated” (fold change >2, FDR <0.001) encoded: 14 SPI-2 proteins; 2 phosphate transport proteins (PtsB and PtsC); two enzymes involved in the arginine biosynthesis pathway (ArgB and ArgC); and the proteins Fis (DNA-binding protein), IolR (repressor of *myo*-inositol utilization), RluD (pseudouridine synthase), WzxE (translocation of the enterobacterial common antigen to the outer membrane), and OmpR. We searched for genes that had not been previously been reported to play a role in virulence in *S*. Typhimurium ST19 [49, 50] and found only three: a SPI-2 gene (*sscB*) and *argBC*.

**Figure 5.**
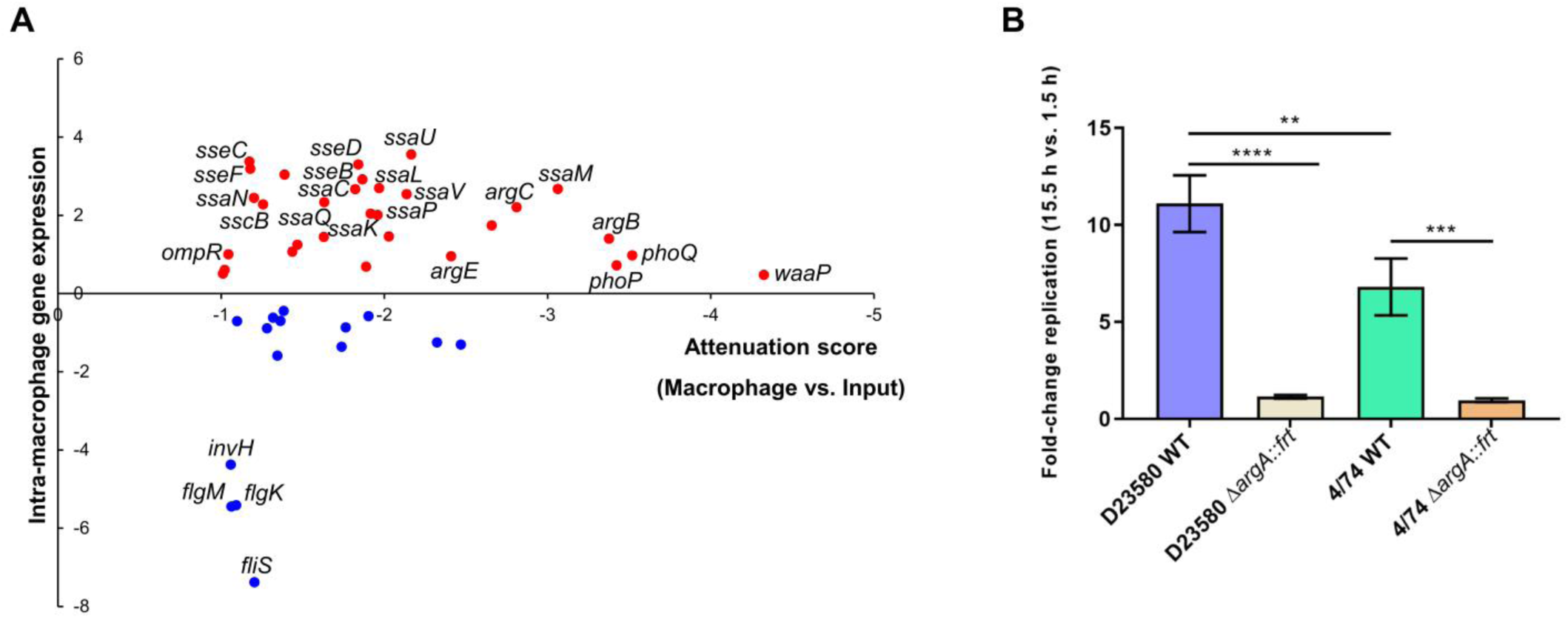
Macrophage-specific genes of S. Typhimurium D23580 are required for virulence in animal infection models. (A) Representation of the 45 genes, among the 68 macrophage-specific genes, in S. Typhimurium D23580 that showed an RNA-seq fold-change with FDR ≤0.05. The RNA-seq fold-change was calculated comparing transcriptomic data from D23580 recovered from the intra-macrophage environment (8 h post-infection) versus D23580 grown to ESP in LB (extracted from Canals and colleagues [15]). The attenuation score represents log2 fold-change of the TIS data obtained comparing the two output Macrophage samples versus the two Input samples. (B) Fold-change replication (15.5 h versus 1.5 h) in murine RAW264.7 macrophages. Average of three independent biological replicates each. Error bars show standard deviation. ****, *P*-value <0.0001; ***, *P*-value = 0.0006; **, *P*-value = 0.0043.

To determine if the requirement for the arginine biosynthetic pathway during intra-macrophage replication was a specific feature of the D23580 strain, mutants in the *argA* gene were constructed in D23580 and the ST19 strain 4/74. ArgA is the first enzyme for the biosynthesis of L-arginine from L-glutamate, a pathway that also includes the biosynthesis of L-ornithine [51]. Of the nine genes encoding products involved for the L-arginine biosynthesis, four were in the 68 macrophage-specific gene list: *argA*, *argCB*, *argE*. Furthermore, the encoded products are also involved in the L-ornithine biosynthetic sub-pathway. The individual Δ*argA*::*frt* mutants of both strains, D23580 and 4/74, showed reduced intra-macrophage replication (Figure 5B). The importance of ArgA for growth inside J774 macrophages has already reported for the *S*. Typhimurium ST19 isolate 14028 [52]. Our results indicate that the requirement for arginine genes inside murine macrophages is not a distinguishing feature of D23580 because the same genes were also required by the ST19 isolate 4/74. The decreased ability of the Δ*argA*::*frt* mutants to proliferate is consistent with previous studies, and suggests that arginine is a limiting factor for *S*. Typhimurium growth inside murine macrophages. The requirement for ArgA for optimal intra-macrophage replication of D23580 validates our TIS-based approach in this infection model.

Taken together, the 68 macrophage-specific gene list included many genes that encode *S.* Typhimurium ST19 virulence factors, and did not include any D23580-specific genes. We conclude that no novel virulence factors required for intra-macrophage replication of *S.* Typhimurium ST313 were identified in our experiments.

### Intergenic regions and sRNAs important for fitness inside murine macrophages

To identify *S*. Typhimurium D23580 intergenic regions and sRNAs that impact upon fitness inside macrophages but not in growth in the laboratory media LB and InSPI2, transposon insertions in short genomic regions were investigated (Materials and Methods). Transposon insertions in ten intergenic regions caused macrophage-specific attenuation (Figure S4C). Four of them were located in the plasmid regions. In pSLT-BT, the intergenic regions included: *spvB*-*spvA* upstream of *spvB* which encodes an ADP-ribosyltransferase that destabilizes actin polymerization of the host cells [53, 54]; *int-dhfrI* encodes an integrase and a trimethoprim resistance gene cassette in the Tn*21*-like element; and the downstream region of *repA* (called *repA_2* in D23580). In the pBT1 plasmid, the intergenic region was located between two genes encoding hypothetical proteins, *pBT1-0171* and *pBT1-0181*. Assuming disruption of promoter regions of the flanking genes, most of the chromosomal intergenic regions were expected to have an effect in fitness inside macrophages: upstream of *dksA*, upstream of a gene located in a SPI-6 associated region, SPI-2, upstream of *pssA* (phosphatidylserine biosynthesis), and upstream of *rpoB* (DNA-directed RNA polymerase subunit beta). An exception was seen upstream of *rmlB* (biosynthesis of O-antigen), where the effects on fitness were likely due to the polarity effects of the insertion of a strong promoter upstream of this gene. Overall, the phenotype of most of these intergenic transposon insertions was supported by previous studies that showed that the particular downstream genes were important for growth and intracellular survival inside macrophages.

Only transposon insertions in one sRNA caused attenuation in the intra-macrophage environment but not in the LB and InSPI2 *in vitro* growth conditions, namely AmgR (Figure S4D). This sRNA is an antisense RNA of the *mgtC* gene [55]. AmgR attenuates virulence mediated by decreasing MgtC protein levels [55]. The fact that disruptions in this sRNA attenuate D23580 within macrophages should be interpreted with caution because transposon insertions disrupt both DNA strands and AmgR overlaps the *mgtC* gene, which is known to be critical for macrophage survival [56]. This finding may simply reflect the known role of *mgtC* in macrophage infection and not the involvement of AmgR in modulating MtgC protein levels.

### Limitations of this study

As for all global mutagenesis approaches, it is important to consider the limitations of our strategy. First, the relatively large size of the mutant pools generated a highly competitive environment, in which trans-complementation could occur. This phenomenon is characterized by compensating genetic defects in some mutants by the presence of the functional genes in other mutants. Second, because the Tn*5* transposon carries an outward facing promoter that drives expression of the Km resistance gene, individual transposon insertions can cause polar effects due to the increased transcription of downstream genes [57, 58]. Third, in the case of macrophage infection, although it would be ideal if individual macrophages were only infected by a single Tn*5*-carrying bacterium, the final multiplicity of infection (M.O.I.) was on average 42:1, meaning that combinations of mutants could have co-localized within the same intra-macrophage vacuole. The genes that contribute to intra-macrophage fitness that were identified here reflected selection for mutants with defects in the ability to replicate and survive inside macrophages, and also selection for mutants lacking certain SPI-1-associated factors such as InvH [59]. The genes required for optimal intra-macrophage fitness of *S*. Typhimurium sequence type ST313 showed substantial overlap with *S*. Typhimurium ST19 genes previously associated to virulence.

### *S*. Typhimurium D23580 has many plasmid-encoded required genes

*S*. Typhimurium D23580 contains four plasmids: pSLT-BT (∼117 kb), pBT1 (∼84.5 kb), pBT2 (∼2.5 kb), and pBT3 (∼2 kb) [14, 15]. The pSLT-BT and pBT1 plasmids have published annotations that were used to study gene requirements for growth and survival in this study. Two pSLT-BT plasmid genes were designated as required, *parA* and *parB* (Figure 6A, Table S2), and the same genes were also found to be required for pBT1 plasmid maintenance (Figure 6B, Table S2). The requirement of ParA and ParB for effective plasmid partitioning means that transposon insertions in both *parA* and *parB* caused plasmid loss in previous TraDIS studies [60]. The unexpected discovery of 34 more pBT1-encoded genes that were required, for stable maintenance of the plasmid or optimal fitness of *S*. Typhimurium D23580, included the *cysS^pBT1^* gene that has been reported previously [15].

**Figure 6.**
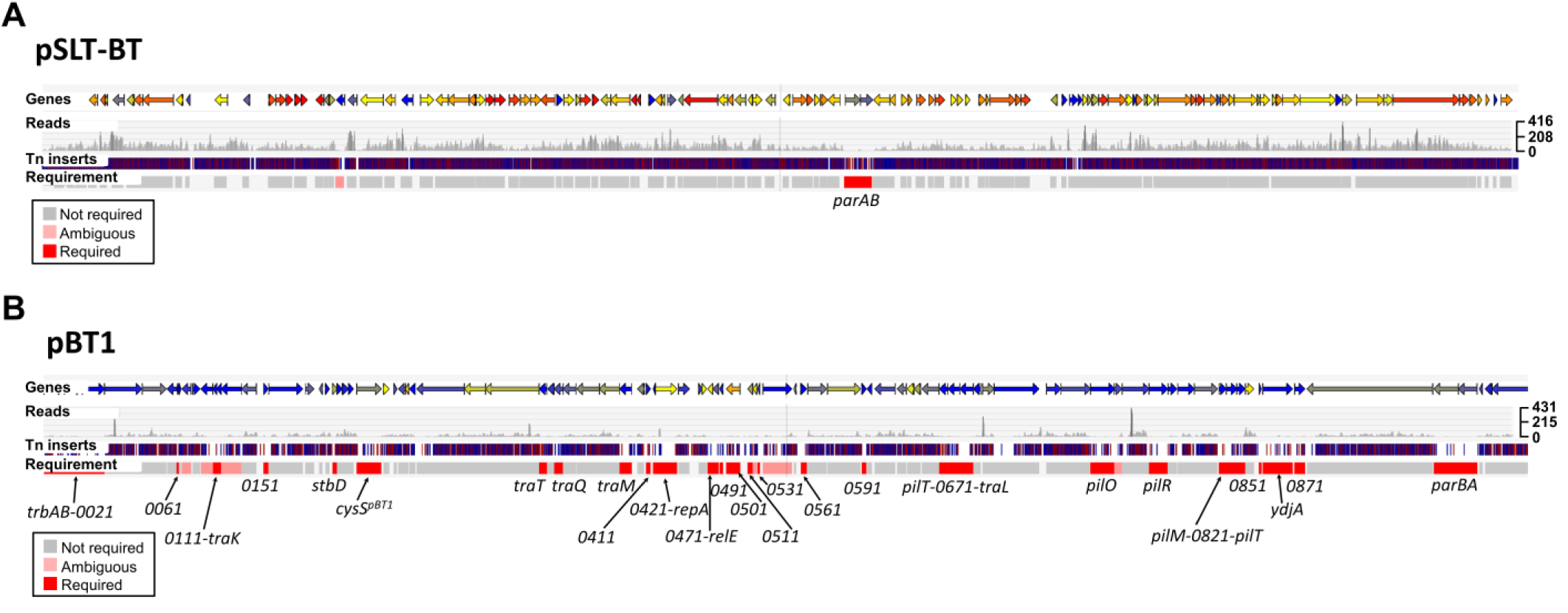
Identification of required genes encoded on S. Typhimurium D23580 pSLT-BT and pBT1 plasmids. (A) pSLT-BT plasmid; and (B) pBT1 plasmid. Figures were obtained using the Dalliance genome viewer (https://hactar.shef.ac.uk/D23580). Coloured arrows at the top represent genes (colour is based on GC content, blue = low, yellow = intermediate, red = high). Each sample is represented by three tracks, in this case the Input is the only sample shown. The first track shows raw data for the Illumina sequencing reads. The second track contains blue and red lines that correspond to transposon insertion sites; red = + orientation of the transposon, same as genes encoded on the plus strand, blue = opposite direction. The third track highlights in red those genes that were considered required for growth in that condition based on an insertion index (Materials and Methods). The names of required genes are indicated at the bottom. The scale on the right represents sequence read coverage.

The pBT1 plasmid is dispensable for *in vitro* growth of *S*. Typhimurium D23580 in LB (Figure 7A, Tables S3 and S5) and required for optimal intra-macrophage replication (Text S1, Figure S6) [15]. The fact that our TIS analysis identified so many pBT1-encoded genes that were required suggests the involvement of multiple genes in the maintenance and replication of the plasmid [60, 61]. For example, the *repA^pBT1^* gene, involved in plasmid replication [62], was required for pBT1 in D23580. The high proportion of pBT1 genes (38%) designated required could reflect the presence of particular features in this plasmid, such as toxin/antitoxin systems and/or DNA-binding proteins. We noted that the percentage of AT content of the pBT1 plasmid is particularly high, 56.7%, compared to the *S*. Typhimurium D23580 chromosome and the pSLT-BT plasmid, which are 47.8% AT and 46.5% AT, respectively. The fact that pBT1 is so AT-rich parallels the AT content of the pSf-R27 plasmid of *Shigella flexneri* 2a strain 2457T which was 55%. An H-NS paralogue, Sfh, is encoded by pSf-R27 and is responsible for a stealth function that allows the plasmid to be transmitted to new bacterial hosts with minimal effects on fitness [63, 64]. Consequently, the introduction of a modified version of the plasmid that lacked *sfh* (pSf-R27Δ*sfh*) into *S*. Typhimurium ST19 SL1344 significantly decreased fitness due to interference with the H-NS regulatory network, whereas the wild-type pSf-R27 plasmid itself did not impact upon fitness when introduced into the same strain [65]. This parallel between the pBT1 and pSf-R27 plasmids raises the possibility that pBT1 encodes an H-NS-like protein that has a global impact upon fitness of D23580, but could not be found by sequence identity alone.

**Figure 7.**
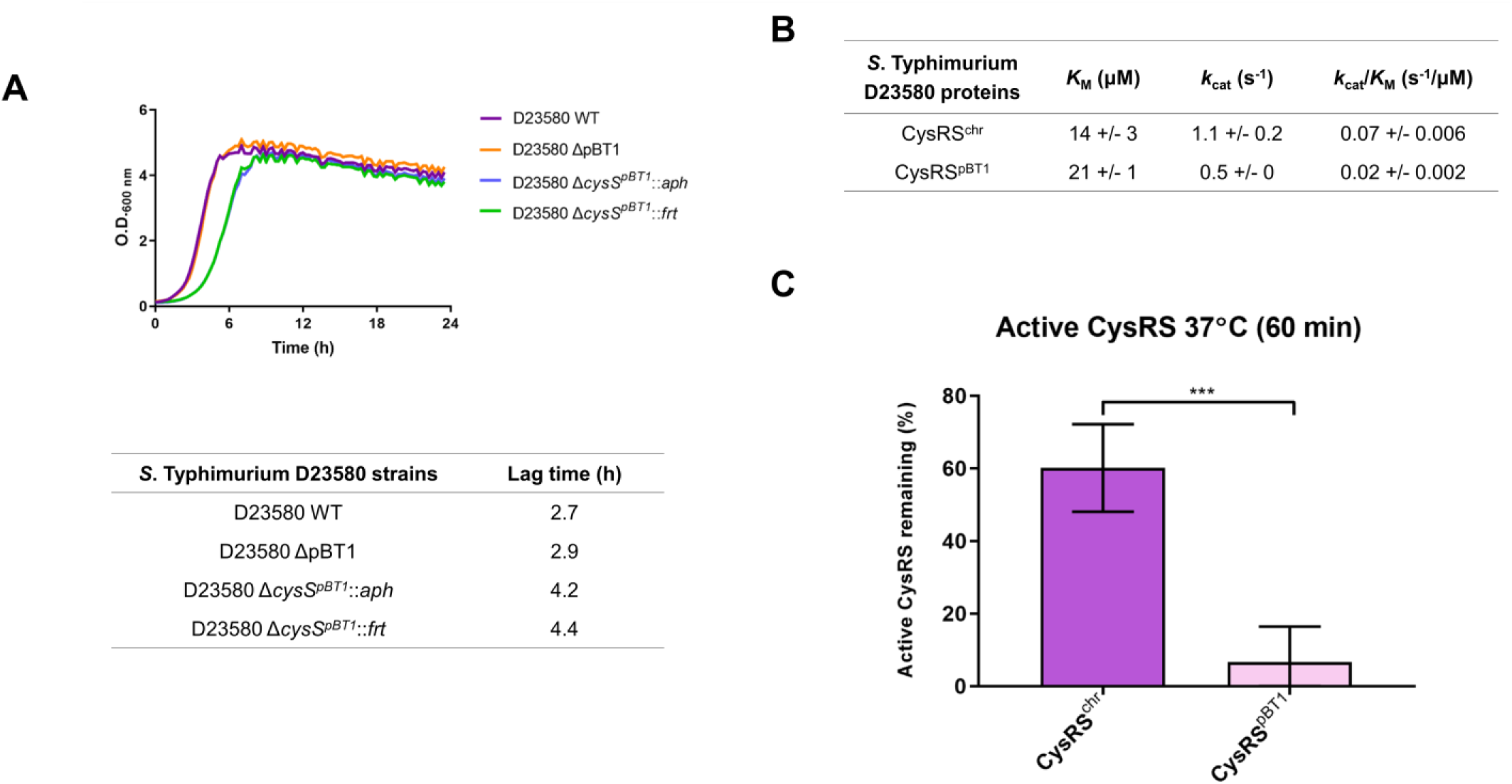
The pBT1 plasmid-encoded CysRS^pBT1^ is less efficient and stable than the chromosomal-encoded CysRS^chr^. (A) Growth curves in LB medium of the D23580 WT strain, the pBT1-cured D23580 strain, and the Km resistant (*aph*) and deletion versions (*frt*) of the *cysS^pBT1^* mutant. The table highlights differences in the lag time between strains. (B) The catalytic efficiency (*k*cat/*K*M) of CysRS^pBT1^ is 3-fold lower than for CysRS^chr^. Recombinant CysRS^pBT1^ and CysRS^chr^ were purified by overexpression in *E. coli*. ATP-PPi exchange was used to determine the steady-state kinetic parameters for activation of cysteine. (C) CysRS^pBT1^ is 10-fold less stable than CysRS^chr^. This comparison was performed by incubating the enzymes at 37°C for 60 minutes.

To investigate pBT1-specific features that could explain the high number of genes with lower amount of transposon insertions than the average amount in the chromosomal and the pSLT-BT plasmid, genes with annotated known functions were studied. The pBT1 plasmid carries a set of conjugative genes and was successfully introduced into 4/74 by conjugation demonstrating the functionality of those genes. At least 11, out of the 36 required genes in pBT1, encoded putative conjugal proteins that are critical for plasmid transfer (*trbAB*, *traK*, *traT*, *traQ*, *traM*, *pilT*, *traL*, *pilO*, *pilR*, *pilM*, *pilT*) [66]. The transposon-induced overexpression of the *traAB* genes in IncI plasmids has been reported to cause a growth defect [67]. Additionally, the plasmid contains at least one annotated toxin/antitoxin system: *stbD* (*pBT1-0201*)/ *stbE* (*pBT1-0211*). The gene encoding the antitoxin StbD was required, while the gene encoding the toxin StbE was dispensable (Table S2). Two other genes, *relE* (*pBT1-0481*) and *relB* (*pBT1-0521*), are annotated as a toxin and an antitoxin, respectively, and are separated by three genes. Our data showed that the gene encoding the toxin RelE was required, while the gene encoding the antitoxin RelB gave an ambiguous result. The precise role of pBT1-encoded toxin/antitoxin systems upon plasmid biology and the fitness of D23580 merits further investigation.

One of the limitations of TIS approaches is that polarity effects can be caused by the introduction of a strong transposon-encoded promoter in a specific genetic context. Such polarity effects could explain the low density of transposons observed in some regions of the pBT1 plasmid, compared with the pSLT-BT plasmid and other plasmids studied in previous TIS studies. Most of the pBT1-encoded genes are hypothetical, and lack a known function. It remains to be determined if the high expression of certain plasmid regions could be toxic to bacterial cells.

The finding that curing of pBT1 does not impact upon fitness shows that the plasmid is not in itself essential. Previously, the observation of fitness defects associated with mutations in specific plasmid-encoded genes reflected the essentiality of the entire plasmid [68, 69], which is extremely uncommon [70]. Because the fitness defect of the D23580 Δ*cysS^pBT1^* mutant could reflect an uncharacterized aspect of pBT1 biology, it was investigated in more detail.

### The pBT1-encoded CysRS^pBT1^ is less efficient and stable than the chromosomal-encoded CysRS^chr^

Cysteinyl-tRNA synthetase (CysRS) is an essential protein in bacteria involved in translation. Recently, we showed that *S*. Typhimurium D23580 expresses high levels of *cysS^pBT1^* and that *cysS^chr^* is expressed at very low levels [15]. To investigate why an organism would express a plasmid-based gene instead of a chromosomal copy, recombinant CysRS^chr^ and CysRS^pBT1^ were purified by overexpressing the relevant D23580 genes in *Escherichia coli* (*E. coli*) and used for enzymatic analysis. The steady-state kinetic parameters for activation of cysteine by both CysRS^chr^ and CysRS^pBT1^ were determined using ATP-PPi exchange. CysRS^pBT1^ had a 3-fold lower catalytic efficiency (*k*cat/*K*M) than CysRS^chr^ (Figure 7B). Additionally, the stability of the enzymes was determined. CysRS^pBT1^ was 10 times less stable than CysRS^chr^ after incubation at 37°C for 60 min (Figure 7C). The fact that CysRS^pBT1^ was both less efficient at cysteine activation and more unstable over time raises the possibility that *S*. Typhimurium D23580 could be using CysRS^pBT1^ as a trigger to shut down translation during stressful conditions. A less efficient CysRS within the bacterial cell would lead to the accumulation of uncharged tRNAs which could induce the stringent response to help the cell cope with stress [71, 72]. We speculate that the preference for expressing the plasmid-encoded *cysS^pBT1^* could give *S*. Typhimurium D23580 the ability to respond to stress conditions at the level of translation.

To investigate how widespread plasmid-encoded aminoacyl-tRNA synthetases are in biology, a database containing 13,661 bacterial plasmids was generated (Materials and Methods). A total of 79 plasmid gene products were found to contain tRNA synthetase-related functional annotations. Closer inspection of those CDS (coding sequences) revealed two classes: complete aminoacyl-tRNA synthetase CDS (30); and CDS encoding aminoacyl-tRNA synthetase fragments and/or related functions (49) (Table S8). The presence of aminoacyl-tRNA synthetase paralogues and paralogous fragments have been previously reported in eukaryotic and prokaryotic genomes [73]. Our analysis suggests that alternate plasmid-encoded aminoacyl-tRNA synthetases exist, and the molecular function of plasmid-encoded translation-related genes warrants further study.

## Perspective

The significant impact of iNTS disease as a major public health problem in sub-Saharan Africa has led to *S.* Typhimurium ST313 becoming an active focus of research. To understand how African *S.* Typhimurium ST313 causes disease, it was important to determine whether this pathovariant carries novel virulence genes that had not previously been found in *S*. Typhimurium ST19. Our transposon insertion sequencing approach during infection of the murine RAW264.7 macrophage infection model suggests that novel virulence factors are not encoded by *S.* Typhimurium ST313 D23580.

Here, we present an online resource that allows the candidate genes that impact upon fitness of *S*. Typhimurium ST313 strain D23580 to be visualized, both in particular *in vitro* growth conditions and during intra-macrophage replication: https://hactar.shef.ac.uk/D23580.

In terms of plasmid biology, we conclude that, although pBT1 is not essential, this plasmid contains a high proportion of genes that impact upon bacterial fitness: our data hint at a new role for plasmid-encoded aminoacyl-tRNA synthetases.

## Materials and Methods

### Bacterial strains and growth conditions

LB was obtained by mixing 10 g/L tryptone (Difco), 5 g/L yeast extract (Difco), and 5 g/L NaCl (Sigma). The InSPI2 medium was based on PCN (pH 5.8, 0.4 mM Pi), which was prepared as previously described [75]. When required, the antibiotic Km was added to a final concentration of 50 μg/mL, and tetracycline (Tc) to 20 μg/mL.

Bacterial strains used for this study are shown in Table S9. Permission to work with *S*. Typhimurium strain D23580 from Malawi [14] was approved by the Malawian College of Medicine (COMREC ethics no. P.08/4/1614).

### Construction of a transposon library in *S*. Typhimurium D23580

A library of transposon insertion mutants was constructed in *S*. Typhimurium D23580 as previously described with some modifications [20]. Briefly, D23580 was grown in rich medium to logarithmic phase and competent cells were prepared. Transposome mixtures were prepared by mixing the EZ-Tn5 <KAN-2> transposon from Epicentre Biotechnologies, EZ-Tn5 transposase and TypeOne restriction inhibitor, and transposomes were transformed into D23580 competent cells. A total of eight electroporations derived from two transposome mixtures were performed and cells were recovered by addition of SOC medium and incubation at 37°C for 1 h. Bacterial mixtures were plated onto LB agar Km at a concentration of 50 μg/mL and incubated at 37°C overnight. The transposon mutants were collected from the plates by adding LB and were joined to grow them together in LB Km 50 μg/mL at 37°C overnight.

### Passages of the *S*. Typhimurium D23580 transposon library in LB and InSPI2

The D23580 transposon library was grown in LB Km 50 μg/mL at 37°C 220 rpm for 16 h, and genomic DNA was purified from a fraction of the bacterial culture (input). Another fraction of the bacterial culture was washed twice with PBS and resuspended in LB or InSPI2 media. A dilution 1:100 (∼2.5 x 10^8^ cells) was inoculated into 25 mL of LB or InSPI2 media (without antibiotic), respectively, and cultures were incubated at 37°C, 220 rpm for 24 h (passage 1). Two more passages were performed in an InSPI2 media. For LB passages, 250 μL were transferred in each individual passage, after two washes with PBS (∼1.4 x 10^9^ cells). For InSPI2 passages, 640 μL were transferred into the next passage (∼1.7 x 10^8^ cells). Genomic DNA was purified from the third passage in LB (output LB) and InSPI2 (output InSPI2). Genomic DNA purifications were performed using the DNeasy Blood & Tissue Kit (Qiagen) following manufacturer’s indications for extractions from gram-negative bacteria.

### Library preparation and Illumina sequencing for the input and the LB and InSPI2 output samples

Genomic DNAs were fragmented to 300 bp with a S220/E220 focused-ultrasonicator (Covaris). Samples were prepared using the NEBNext DNA Library Prep Master Mix Set for Illumina for use with End User Supplied Primers and Adapters (New England Biolabs) following manufacturer’s instructions. The DNA fragments were end-repaired and an “A” base was added to the 3’ends prior to ligation of the Illumina adapters (*PE Adapters* from Illumina). In order to amplify the transposon-flanking regions, transposon-specific forward oligonucleotides were designed such that the first 10 bases of each read would be the transposon sequence: PE PCR Tn-12, for the input sample (Input); PE PCR Tn-1 and PE PCR Tn-7, for the output sample in LB; and PE PCR Tn-10 and PE PCR Tn-4, for the output sample in InSPI2 (S10 Table). These oligonucleotides were used for PCR amplification with the Illumina reverse primer *PE PCR Primer 2.0*. The primers included the adapters and the sequences necessary for attachment to the Illumina flow cell. Furthermore, a specific 6-base barcode included in the forward primer was incorporated into each of the samples in order to pool them together in a single lane for sequencing. Oligonucleotides ordered were HPLC-purified with a phosphorothioate bond at the 3’end from Eurofins Genomics. The three libraries of amplified products were pooled in equimolar amounts and size-selected to 200-500 bp. After QC assessment, the pool was paired-end sequenced, using the Illumina sequencing primers, in one lane on a HiSeq 2500 at 2х125 bp. 15% of library of the bacteriophage ΦX174 genome, provided by Illumina as a control, was added to the lane to overcome the low complexity of the bases after the barcode in Read 1.

### Sequence analysis of the *S*. Typhimurium D23580 transposon library

Cutadapt version 1.8.1 was used to demultiplex the sequence reads based on the 6-base barcode [76]. Transposon sequences at the beginning of the reads were removed using the same program. BWA-MEM [77] was used to map the reads against the *S*. Typhimurium D23580 genome sequence (accession: PRJEB28511). Reads with a mapping quality <10 were discarded, as were alignments which did not match at the 5’end of the read (immediately adjacent to the transposon) (Table S1). The exact position of the transposon insertion sites and the frequency of transposons for every annotated gene were determined.

An insertion index was calculated for each gene as explained in [19], using the Bio::Tradis toolkit [78]. Genes with insertion index values <0.05 (cut-off determined using Bio::Tradis) were considered as “required” for growth in the Lennox rich medium. Genes with an insertion index between 0.05 and 0.075 were considered “ambiguous”, and genes with an insertion index >0.075 were considered “not required”. Table S2 shows the number of reads, transposon insertion sites, insertion index, and essentiality call per gene.

### Infection of RAW264.7 macrophages with *S*. Typhimurium D23580

The D23580 transposon library was grown in LB Km 50 μg/mL at 37°C 220 rpm for 16 h, and genomic DNA was purified from two different biological replicates as input samples. Murine RAW264.7 macrophages (ATCC TIB-71) were grown in DMEM high glucose (Thermo Fisher Scientific) supplemented with 10% heat-inactivated fetal bovine serum (Thermo Fisher Scientific), 1X MEM non-essential amino acids (Thermo Fisher Scientific) and 2 mM L-glutamine (Thermo Fisher Scientific), at 37°C in a 5% CO_2_ atmosphere. 10^6^ macrophage cells were seeded on each well of 6-well plates (Sarstedt) 24 h before infection. Bacteria were opsonized with 10% BALB/c mouse serum (Charles River) in 10 volumes of DMEM for 30 min on ice. Macrophages were infected with *Salmonella* at an M.O.I. of approximately 10:1, and infections were synchronized by centrifugation (5 min at 1,000 rpm). After 30 min of infection, macrophages were washed with DPBS (Thermo Fisher Scientific), and DMEM with supplements and gentamicin 100 μg/mL was added to kill extracellular bacteria. After 1 h, macrophages were washed with DPBS, and fresh DMEM with supplements and gentamicin 10 μg/mL was provided for the rest of the incubation time at 37°C with 5% CO_2_. In some wells, 1% Triton X-100 was added to recover intracellular bacteria and plate dilutions to obtain bacterial counts for the 1.5 h time point. For the rest of the wells, after 12 h from the initial infection, macrophages were washed with DPBS and intracellular bacteria were collected using 1% Triton X-100. Some wells were used for obtaining bacterial counts and calculate the fold-change replication of the intracellular bacteria (12 h versus 1.5 h) (Figure S3B). For the output samples of the D23580 transposon library, 12 wells (of 6-well plates) were pooled for each replicate. The samples of 1% Triton X-100 containing the intracellular bacteria were centrifugated and the supernatants were discarded. Pellets were resuspended in LB and transferred into a flask to grow bacteria for 10 h 220 rpm in LB supplemented with Km. Genomic DNA was extracted from those cultures and prepared for Illumina sequencing, together with the input samples.

For macrophage infections with the individual *S*. Typhimurium strains D23580 WT, D23580 Δ*argA*::*frt*, 4/74 WT, and 4/74 Δ*argA*::*frt*, intracellular bacteria were recovered after 1.5 h and 15.5 h. Bacterial counts at the two time points were used to calculate the fold-change replication in the intra-macrophage environment (Figure 5B).

### Library preparation and Illumina sequencing for the input and output samples of the RAW264.7 macrophage experiment

Genomic DNA samples were prepared for Illumina sequencing following the previously described protocol. The transposon-specific forward oligonucleotides used for amplifying the sequence adjacent to the transposon were: PE PCR Tn-12, for the input biological replicate 1 sample (Input 1); PE PCR Tn-7, for the input biological replicate 2 sample (Input 2); PE PCR Tn-1, for the output biological replicate 1 sample (Macrophage 1); and PE PCR Tn-5, for the output biological replicate 2 sample (Macrophage 2) (Table S10). In this case, oligonucleotides were ordered HPLC-purified with a phosphorothioate bond at the 3’end from Integrated DNA Technologies (IDT).

The four libraries of amplified products were pooled in equimolar amounts with two other libraries and size-selected and QC-assessed. The pool was paired-end sequenced, using the Illumina sequencing primers, in one lane on a HiSeq 4000 at 2х150 bp. In this case, 50% of a library of the bacteriophage ΦX174 genome was added to the lane to overcome the low complexity of the bases after the barcode in Read 1.

### Sequence analysis of the *S*. Typhimurium D23580 transposon library

Analysis was performed using DESeq2 and following the same strategy described in [79]. Results are shown in log2 fold-change. A cutoff of 2-fold-change and *P*-value ≤0.05 was applied (Table S7).

### Construction of mutants in *S*. Typhimurium D23580 and 4/74 by λ Red recombineering

Mutants were constructed using the λ Red recombination system [80]. Using oligonucleotides Fw-argA-P1 and Rv-argA-P2, a Km resistance cassette was PCR-amplified from plasmid pKD4. The PCR product was transformed by electroporation into D23580 and 4/74 containing the pSIM5-*tet* plasmid to replace the *argA* gene. Recombinants were selected on LB agar plates supplemented with Km. The 4/74 Δ*argA*::*aph* construction was transduced into WT 4/74 using the generalized transducing bacteriophage P22 HT105/1 *int-201* as previously described [27]. The antibiotic resistance cassettes from both, the 4/74 Δ*argA*::*aph* and D23580 Δ*argA*::*aph*, were removed by the use of the pCP20-TcR plasmid [81]. Strains and plasmids are included in Table S9 and oligonucleotides in Table S10.

### CysRS cloning and purification

Genomic DNA from *S.* Typhimurium D23580 was used to PCR amplify both chromosomal and plasmid *cysS* which were then cloned into pET28a(+). Chromosomal and plasmid CysRS were expressed in *E. coli* BL21(DE3) with 1 mmol IPTG induction for 4 h. Cells were harvested, lysed by sonication and purified using a TALON metal affinity resin. CysRS was eluted with 250 mM imidazole and fractions containing protein were concentrated and dialyzed overnight in 50 mM Tris pH 7.5, 100 mM KCl, 5 mM MgCl_2_x, 3 mM 2-mercaptoethanol, 5% glycerol, and then dialyzed 4 h in similar buffer with 50% glycerol for storage. Proteins were stored at −20°C. Oligonucleotides used for cloning and expression are included in Table S10.

### CysRS pyrophosphate exchange – steady state kinetics

To determine the *K*_M_ for Cys, pyrophosphate exchange was completed in a reaction containing 100 mM HEPES pH 7.5, 30 mM KCl, 10 mM MgCl_2_ 1 mM NaF, 25 nM CysRS, 50 µM-2mM Cys, 2 mM ATP, 2 mM ^32^P-PPi. The reaction without CysRS was incubated at 37°C for 5 min at which point the enzyme was added. Then aliquots were taken at 1-4 min by combining the reaction with quench solution (1% activated charcoal, 5.6% HClO_4_, 1.25 M PPi). On a vacuum filter with 3 mm filter discs, filter discs were pre-rinsed with water, charcoal reaction added, washed 3x H_2_O and 1x 95% EtOH. Radiation was quantified using liquid scintillation counting. Michaelis-Menton equation was used to determine kinetic parameters.

### CysRS thermal stability

To determine the stability of protein, chromosomal and pBT1 CysRS were incubated at 37°C for 0 and 60 min and active site titration was used to measure the activity of the protein. Active site titration was completed in 30 mM KCl, 10 mM MgCl_2_, 80uM ^35^S-Cys, 2 mM ATP, pyrophosphatase and CysRS. At 0 min, 5 μL of CysRS were added to the reaction mixture and placed at 37°C for 10 min. The reaction was quenched by placing tubes on ice. After 60 min incubation of purified protein at 37°C, 5 μL were combined with the reaction mixture as above. All reactions were placed on a vacuum filtration unit on a Protran BA85 nitrocellulose membrane, washed three times with 1 mL 15 mM KCl and 5 mM MgCl_2_ and dried. Then 4 mL of liquid scintillation cocktail were added and radiation was quantified using liquid scintillation counting.

### Growth curves of the D23580 bacterial strains

To determine the growth rate of the D23580 WT, D23580 ΔpBT1, D23580 Δ*cysS^pBT1^*::*aph* and D23580 Δ*cysS^pBT1^*::*frt* strains in LB, a Growth Profiler 960 was used (EnzyScreen). Bacterial cells grown for 16 h in LB, 37°C 220 rpm, were diluted in 250 μl of LB to OD600 0.01 and incubated in the Growth Profiler for 24 h at 37°C, shaking at 224 rpm. The OD600 values were measured every 15 min.

### Statistical analyses

Graphpad Prism 8.0.1 was used for statistical analyses (GraphPad Software Inc., La Jolla, CA, USA). One-way ANOVA and Tukey’s multiple comparison test were used for comparative analyses.

### Analysis of conservation of aminoacyl-tRNA synthetases in bacterial plasmids

A plasmid database was generated from the plasmid sub-section of NCBI’s Reference Sequence Database (RefSeq) v90 [82], and included 13,924 complete plasmid sequences. The associated taxonomical information was downloaded and non-bacterial sequences were removed to leave 13,661 plasmids. The GenPept files were processed to extract the associated coding sequence annotations.

## Acknowledgments

We are thankful to present and former members of the Hinton laboratory for helpful discussions, particularly Xiaojun Zhu and Carsten Kröger, and to Paul Loughnane for his technical assistance. We are grateful to John Kenny for his expertise in library preparation for Illumina sequencing, and thank the Centre for Genomic Research at the University of Liverpool for the use of the Covaris equipment. We appreciated our helpful discussions with Duy Phan concerning the design and analysis of the transposon insertion experiments.

## Supporting information

**Table S1. Number of sequenced reads for each sample at every step**. Only R1 reads are included. The percentages were calculated relative to the previous step with the exception of deduplication, which was calculated relative to the number of mapped reads.

**Table S2. Number of reads, transposon insertion sites, insertion index, and essentiality call per gene**. Samples included: Input (for LB and InSPI2), LB (output), InSPI2 (output), Input 1 (for Macrophage), Input 2 (for Macrophage), Macrophage 1 (output), and Macrophage 2 (output).

**Table S3. Raw data for figures.**

**Figure S1. Identification of genes that are required in *S*. Typhimurium D23580 but not in other *Salmonella* pathovariants.** (A) Comparative analysis of *S*. Typhimurium D23580 required genes with previously identified required genes in TIS studies in *S*. Typhimurium [20, 24], and *S*. Typhi [19, 20]. SL3261 was derived from SL1344; and WT174 was derived from Ty2. For the previously published studies, only genes that shared an ortholog in D23580 were included for the analysis. Individual Venn diagram analyses including a comparison with only *S*. Typhimurium (B) and *S*. Typhi (C) strains were also generated.

**Table S4.** Raw data for supporting figures.

**Figure S2. *S*. Typhimurium D23580 genes with reported H-NS binding sites in *S*. Typhimurium SL1344.** Individual growth curves, in LB medium, of the Km resistant versions (*aph*, *n* = 8) and the deletion versions (*frt*, *n* = 7) of (A) a SPI-1 mutant (*hilC*), a SPI-2 mutant (*ssrAB*), and (B) LPS mutants (*waaL* and *waaG*).

**Table S5. Lag time, maximum OD_600_ and maximal growth rate from growth curves for different *S*. Typhimurium D23580 mutants and the WT strain.**

**Table S6. Number of reads, transposon insertion sites, insertion index, and essentiality call per intergenic regions ≥100 bp and sRNAs.**

**Figure S3. *S*. Typhimurium D23580 transposon library passaged three times in murine RAW264.7 macrophages.** (A) M.O.I. (number of bacterial cells used to infect 1 macrophage) of the D23580 transposon library infecting murine RAW264.7 macrophages for 8 h, used in the first (*n* = 1), second (*n* = 3) and third infections (*n* = 3). Fold-change replication of the intra-macrophage bacteria (8 h versus 1.5 h) of the D23580 transposon library seen after each passage. Error bars show standard deviation (*n* = 3). (B) M.O.I. and fold-change replication of the intra-macrophage bacteria of the D23580 transposon library at 12 h p.i. (C) Fold-change replication of the D23580 WT and 4/74 WT strains inside murine RAW264.7 macrophages. M.O.I.s calculated for one of the three biological replicates are indicated at the top of each bar (*n* = 3). (D) The percentage of rough mutants increased after passages of *S*. Typhimurium D23580 WT in macrophages and LB (first and second infections, *n* = 1; third infection, *n* = 3; after third passage, *n* = 3); (E) and in *S*. Typhimurium 4/74 WT (first and second infections, *n* = 1; third infection, *n* = 2; after third passage, *n* = 3).

**Text S1. Supporting results and methods.**

**Figure S4. 206 *S*. Typhimurium D23580 genes cause fitness alteration in macrophages.** (A) 10% of the *S*. Typhimurium D23580 genes are required for growth in all three LB input mutant pools. (B) Identification of *S*. Typhimurium D23580 “macrophage-specific” and “macrophage-associated” genes. The Venn diagram compares the 206 *S*. Typhimurium D23580 genes that showed attenuation in RAW264.7 macrophages when disrupted by a transposon insertion with required genes in the three inputs (Inputs), and the LB and InSPI2 outputs. (C) Venn diagrams including only intergenic regions, and (D) sRNAs.

**Table S7. Analysis of the TIS macrophage data.** Read counts for the two inputs and two outputs (Macrophage), and log2 fold-changes and adjusted *P*-values for the comparative analysis of each coding gene, noncoding sRNA, and intergenic regions in *S*. Typhimurium D23580.

**Figure S5. Macrophage-attenuated genes of D23580 required for virulence of *S*. Typhimurium in other infection models.** (A) 63% of the D23580 macrophage-attenuated genes are important for virulence of *S*. Typhimurium 4/74 in food-related animal infection models [49]. Only orthologous chromosomal and pSLT plasmid genes were included for the analysis. (B) D23580 macrophage-attenuated genes compared to *S*. Typhimurium 14028 genes associated to virulence in BALB/c mice [50]. Only orthologous chromosomal genes were included for the analysis.

**Figure S6.** The *S*. Typhimurium D23580 Δ*STM2475* mutant and the pBT1-cured strain exhibited decreased proliferation in the intra-macrophage environment. (A) Intra-macrophage proliferation assays of the D23580 WT, D23580 Δ*STM2475*::*frt*, D23580 *STM2475*^4/74SNP^, D23580 ΔpBT1; and 4/74 WT, 4/74 Δ*STM2475*::*frt*, 4/74 *STM2475*^D23580SNP^, 4/74 Δ*ssrAB*::*frt*. Bars represent average of three independent biological replicates and standard deviation. Significant differences indicate *P-value*: ***, 0.0002; **, 0.0011; *, 0.0116; ns, not significant. (B) Alignment of the *STM2475* promoter region in four *S.* Typhimurium strains. (C) Conservation of the nucleotide indel in *S.* Typhimurium ST313 strains.

**Figure S7. The *STM1630* gene is inactivated in *S.* Typhimurium D23580**. (A) Transposon insertion profile of the *STM1630* region from our D23580 Dalliance genome browser. (B) There were not significant differences in growth in LB between D23580 WT and the *ΔSTM1630* mutants, with or without the Km resistance cassette. (C) Absolute expression levels of *STM1630* in *S.* Typhimurium D23580 and 4/74 (extracted from Canals and colleagues [15]). Values represent TPM, TPM ≤ 10 means no expression. (D) Disruption of STM1630 − 10 box in the promoter region: two SNP-difference between 4/74 and D23580. (E) The D23580 isoform is conserved in all ST313 genomes analyzed, including lineage 1 and 2 and UK-ST313 strains described in Ashton and colleagues [83]. BLASTn was used to identify the genotype of the *STM1630* transcriptional start site −10 region in all genomes and the results were visualized in the context of the phylogenetic tree from Ashton and colleagues [83]. (F) Intra-macrophage proliferation assays of the D23580 and 4/74 WT strains, the *ΔSTM1630::frt* mutants for D23580 and 4/74, and the D23580 *STM1630^4/74SNP^* mutant. Bars represent average of three independent biological replicates and standard deviation. Significant differences indicate *P*-value: ****, <0.0001; ***, <0.001; ns, not significant.

**Table S8. 79 aminoacyl-tRNA synthetases found in the custom bacterial plasmid database.**

**Table S9. Bacterial strains used in this study.**

**Table S10. Oligonucleotides used in this study.**

## References

1. Majowicz SE, Musto J, Scallan E, Angulo FJ, Kirk M, O’Brien SJ, et al. The Global Burden of Nontyphoidal Salmonella Gastroenteritis. Clin Infect Dis. 2010;50: 882–889. doi:10.1086/650733

2. Stecher B, Robbiani R, Walker AW, Westendorf AM, Barthel M, Kremer M, et al. Salmonella enterica Serovar Typhimurium Exploits Inflammation to Compete with the Intestinal Microbiota. PLOS Biol. 2007;5: e244. doi:10.1371/journal.pbio.0050244

3. Winter SE, Thiennimitr P, Winter MG, Butler BP, Huseby DL, Crawford RW, et al. Gut inflammation provides a respiratory electron acceptor for Salmonella. Nature. 2010;467: 426–429. doi:10.1038/nature09415

4. Fields PI, Swanson RV, Haidaris CG, Heffron F. Mutants of Salmonella typhimurium that cannot survive within the macrophage are avirulent. Proc Natl Acad Sci. 1986;83: 5189–5193. doi:10.1073/pnas.83.14.5189

5. Feasey NA, Dougan G, Kingsley RA, Heyderman RS, Gordon MA. Invasive non-typhoidal salmonella disease: an emerging and neglected tropical disease in Africa. The Lancet. 2012;379: 2489–2499. doi:10.1016/S0140-6736(11)61752-2

6. Gordon M, Banda H, Gondwe M, Gordon S, Boeree M, Walsh A, et al. Non-typhoidal salmonella bacteraemia among HIV-infected Malawian adults: high mortality and frequent recrudescence. Aids. 2002;16: 1633–1641.

7. MacLennan CA, Gilchrist JJ, Gordon MA, Cunningham AF, Cobbold M, Goodall M, et al. Dysregulated Humoral Immunity to Nontyphoidal Salmonella in HIV-Infected African Adults. Science. 2010;328: 508–512. doi:10.1126/science.1180346

8. Raffatellu M, Santos RL, Verhoeven DE, George MD, Wilson RP, Winter SE, et al. Simian immunodeficiency virus–induced mucosal interleukin-17 deficiency promotes Salmonella dissemination from the gut. Nat Med. 2008;14: 421–428. doi:10.1038/nm1743

9. Lê-Bury G, Niedergang F. Defective Phagocytic Properties of HIV-Infected Macrophages: How Might They Be Implicated in the Development of Invasive Salmonella Typhimurium? Front Immunol. 2018;9. doi:10.3389/fimmu.2018.00531

10. Kariuki S, Revathi G, Kariuki N, Kiiru J, Mwituria J, Muyodi J, et al. Invasive multidrug-resistant non-typhoidal Salmonella infections in Africa: zoonotic or anthroponotic transmission? J Med Microbiol. 2006;55: 585–591. doi:10.1099/jmm.0.46375-0

11. Gordon MA, Graham SM, Walsh AL, Wilson L, Phiri A, Molyneux E, et al. Epidemics of Invasive Salmonella enterica Serovar Enteritidis and S. enterica Serovar Typhimurium Infection Associated with Multidrug Resistance among Adults and Children in Malawi. Clin Infect Dis. 2008;46: 963–969. doi:10.1086/529146

12. Su L-H, Chiu C-H, Chu C, Ou JT. Antimicrobial Resistance in Nontyphoid Salmonella Serotypes: A Global Challenge. Clin Infect Dis. 2004;39: 546–551. doi:10.1086/422726

13. Ao TT, Feasey NA, Gordon MA, Keddy KH, Angulo FJ, Crump JA. Global Burden of Invasive Nontyphoidal Salmonella Disease, 20101. Emerg Infect Dis. 2015;21: 941–949. doi:10.3201/eid2106.140999

14. Kingsley RA, Msefula CL, Thomson NR, Kariuki S, Holt KE, Gordon MA, et al. Epidemic multiple drug resistant Salmonella Typhimurium causing invasive disease in sub-Saharan Africa have a distinct genotype. Genome Res. 2009;19: 2279–2287. doi:10.1101/gr.091017.109

15. Canals R, Hammarlöf DL, Kröger C, Owen SV, Fong WY, Lacharme-Lora L, et al. Adding function to the genome of African Salmonella Typhimurium ST313 strain D23580. PLOS Biol. 2019;17: e3000059. doi:10.1371/journal.pbio.3000059

16. Lokken KL, Walker GT, Tsolis RM. Disseminated infections with antibiotic-resistant non-typhoidal Salmonella strains: contributions of host and pathogen factors. Pathog Dis. 2016;74. doi:10.1093/femspd/ftw103

17. van Opijnen T, Camilli A. Transposon insertion sequencing: a new tool for systems-level analysis of microorganisms. Nat Rev Microbiol. 2013;11. doi:10.1038/nrmicro3033

18. Chao MC, Abel S, Davis BM, Waldor MK. The Design and Analysis of Transposon-Insertion Sequencing Experiments. Nat Rev Microbiol. 2016;14: 119–128. doi:10.1038/nrmicro.2015.7

19. Langridge GC, Phan M-D, Turner DJ, Perkins TT, Parts L, Haase J, et al. Simultaneous assay of every Salmonella Typhi gene using one million transposon mutants. Genome Res. 2009;19: 2308–2316. doi:10.1101/gr.097097.109

20. Canals R, Xia X-Q, Fronick C, Clifton SW, Ahmer BM, Andrews-Polymenis HL, et al. High-throughput comparison of gene fitness among related bacteria. BMC Genomics. 2012;13: 212. doi:10.1186/1471-2164-13-212

21. Khatiwara A, Jiang T, Sung S-S, Dawoud T, Kim JN, Bhattacharya D, et al. Genome Scanning for Conditionally Essential Genes in Salmonella enterica Serotype Typhimurium. Appl Env Microbiol. 2012;78: 3098–3107. doi:10.1128/AEM.06865-11

22. Moraes MH de, Desai P, Porwollik S, Canals R, Perez DR, Chu W, et al. Salmonella Persistence in Tomatoes Requires a Distinct Set of Metabolic Functions Identified by Transposon Insertion Sequencing. Appl Environ Microbiol. 2017;83: e03028–16. doi:10.1128/AEM.03028-16

23. Down TA, Piipari M, Hubbard TJP. Dalliance: interactive genome viewing on the web. Bioinformatics. 2011;27: 889–890. doi:10.1093/bioinformatics/btr020

24. Barquist L, Langridge GC, Turner DJ, Phan M-D, Turner AK, Bateman A, et al. A comparison of dense transposon insertion libraries in the Salmonella serovars Typhi and Typhimurium. Nucleic Acids Res. 2013;41: 4549–4564. doi:10.1093/nar/gkt148

25. Knuth K, Niesalla H, Hueck CJ, Fuchs TM. Large-scale identification of essential Salmonella genes by trapping lethal insertions. Mol Microbiol. 51: 1729–1744. doi:10.1046/j.1365-2958.2003.03944.x

26. Thiele I, Hyduke DR, Steeb B, Fankam G, Allen DK, Bazzani S, et al. A community effort towards a knowledge-base and mathematical model of the human pathogen Salmonella Typhimurium LT2. BMC Syst Biol. 2011;5: 8. doi:10.1186/1752-0509-5-8

27. Owen SV, Wenner N, Canals R, Makumi A, Hammarlöf DL, Gordon MA, et al. Characterization of the Prophage Repertoire of African Salmonella Typhimurium ST313 Reveals High Levels of Spontaneous Induction of Novel Phage BTP1. Front Microbiol. 2017;8. doi:10.3389/fmicb.2017.00235

28. Baba T, Ara T, Hasegawa M, Takai Y, Okumura Y, Baba M, et al. Construction of Escherichia coli K-12 in-frame, single-gene knockout mutants: the Keio collection. Mol Syst Biol. 2006;2: 2006.0008. doi:10.1038/msb4100050

29. Lodge JK, Weston-Hafer K, Berg DE. Transposon Tn5 Target Specificity: Preference for Insertion at G/C Pairs. Genetics. 1988;120: 645–650.

30. Goryshin IY, Miller JA, Kil YV, Lanzov VA, Reznikoff WS. Tn5/IS50 target recognition. Proc Natl Acad Sci. 1998;95: 10716–10721.

31. Manna D, Porwollik S, McClelland M, Tan R, Higgins NP. Microarray analysis of Mu transposition in Salmonella enterica, serovar Typhimurium: transposon exclusion by high-density DNA binding proteins. Mol Microbiol. 2007;66: 315–328. doi:10.1111/j.1365-2958.2007.05915.x

32. Manna D, Breier AM, Higgins NP. Microarray analysis of transposition targets in Escherichia coli: The impact of transcription. Proc Natl Acad Sci. 2004;101: 9780–9785. doi:10.1073/pnas.0400745101

33. Lucchini S, Rowley G, Goldberg MD, Hurd D, Harrison M, Hinton JCD. H-NS Mediates the Silencing of Laterally Acquired Genes in Bacteria. PLOS Pathog. 2006;2: e81. doi:10.1371/journal.ppat.0020081

34. Navarre WW, Porwollik S, Wang Y, McClelland M, Rosen H, Libby SJ, et al. Selective Silencing of Foreign DNA with Low GC Content by the H-NS Protein in Salmonella. Science. 2006;313: 236–238. doi:10.1126/science.1128794

35. Kimura S, Hubbard TP, Davis BM, Waldor MK. The Nucleoid Binding Protein H-NS Biases Genome-Wide Transposon Insertion Landscapes. mBio. 2016;7: e01351–16. doi:10.1128/mBio.01351-16

36. Dillon SC, Cameron ADS, Hokamp K, Lucchini S, Hinton JCD, Dorman CJ. Genome-wide analysis of the H-NS and Sfh regulatory networks in Salmonella Typhimurium identifies a plasmid-encoded transcription silencing mechanism. Mol Microbiol. 2010;76: 1250–1265. doi:10.1111/j.1365-2958.2010.07173.x

37. Sturm A, Heinemann M, Arnoldini M, Benecke A, Ackermann M, Benz M, et al. The Cost of Virulence: Retarded Growth of Salmonella Typhimurium Cells Expressing Type III Secretion System 1. PLOS Pathog. 2011;7: e1002143. doi:10.1371/journal.ppat.1002143

38. Ali SS, Soo J, Rao C, Leung AS, Ngai DH-M, Ensminger AW, et al. Silencing by H-NS Potentiated the Evolution of Salmonella. PLoS Pathog. 2014;10. doi:10.1371/journal.ppat.1004500

39. Kong Q, Yang J, Liu Q, Alamuri P, Roland KL, Curtiss R. Effect of Deletion of Genes Involved in Lipopolysaccharide Core and O-Antigen Synthesis on Virulence and Immunogenicity of Salmonella enterica Serovar Typhimurium. Infect Immun. 2011;79: 4227–4239. doi:10.1128/IAI.05398-11

40. Azriel S, Goren A, Rahav G, Gal-Mor O. The Stringent Response Regulator DksA Is Required for Salmonella enterica Serovar Typhimurium Growth in Minimal Medium, Motility, Biofilm Formation, and Intestinal Colonization. Infect Immun. 2015;84: 375–384. doi:10.1128/IAI.01135-15

41. Sittka A, Lucchini S, Papenfort K, Sharma CM, Rolle K, Binnewies TT, et al. Deep Sequencing Analysis of Small Noncoding RNA and mRNA Targets of the Global Post-Transcriptional Regulator, Hfq. PLOS Genet. 2008;4: e1000163. doi:10.1371/journal.pgen.1000163

42. Holmqvist E, Wright PR, Li L, Bischler T, Barquist L, Reinhardt R, et al. Global RNA recognition patterns of post-transcriptional regulators Hfq and CsrA revealed by UV crosslinking in vivo. EMBO J. 2016;35: 991–1011. doi:10.15252/embj.201593360

43. Fröhlich KS, Papenfort K, Berger AA, Vogel J. A conserved RpoS-dependent small RNA controls the synthesis of major porin OmpD. Nucleic Acids Res. 2012;40: 3623–3640. doi:10.1093/nar/gkr1156

44. Fröhlich KS, Haneke K, Papenfort K, Vogel J. The target spectrum of SdsR small RNA in Salmonella. Nucleic Acids Res. 2016;44: 10406–10422. doi:10.1093/nar/gkw632

45. Lévi-Meyrueis C, Monteil V, Sismeiro O, Dillies M-A, Monot M, Jagla B, et al. Expanding the RpoS/σS-Network by RNA Sequencing and Identification of σS-Controlled Small RNAs in Salmonella. PLOS ONE. 2014;9: e96918. doi:10.1371/journal.pone.0096918

46. Hölzer SU, Schlumberger MC, Jäckel D, Hensel M. Effect of the O-Antigen Length of Lipopolysaccharide on the Functions of Type III Secretion Systems in Salmonella enterica. Infect Immun. 2009;77: 5458–5470. doi:10.1128/IAI.00871-09

47. Srikumar S, Kröger C, Hébrard M, Colgan A, Owen SV, Sivasankaran SK, et al. RNA-seq Brings New Insights to the Intra-Macrophage Transcriptome of Salmonella Typhimurium. PLOS Pathog. 2015;11: e1005262. doi:10.1371/journal.ppat.1005262

48. Hammarlöf DL, Canals R, Hinton JC. The FUN of identifying gene function in bacterial pathogens; insights from Salmonella functional genomics. Curr Opin Microbiol. 2013;16: 643–651. doi:10.1016/j.mib.2013.07.009

49. Chaudhuri RR, Morgan E, Peters SE, Pleasance SJ, Hudson DL, Davies HM, et al. Comprehensive Assignment of Roles for Salmonella Typhimurium Genes in Intestinal Colonization of Food-Producing Animals. PLOS Genet. 2013;9: e1003456. doi:10.1371/journal.pgen.1003456

50. Silva-Valenzuela CA, Molina-Quiroz RC, Desai P, Valenzuela C, Porwollik S, Zhao M, et al. Analysis of Two Complementary Single-Gene Deletion Mutant Libraries of Salmonella Typhimurium in Intraperitoneal Infection of BALB/c Mice. Front Microbiol. 2016;6. doi:10.3389/fmicb.2015.01455

51. Charlier D, Glansdorff N. Biosynthesis of Arginine and Polyamines. EcoSal Plus. 2004;1. doi:10.1128/ecosalplus.3.6.1.10

52. Köstner M, Schmidt B, Bertram R, Hillen W. Generating Tetracycline-Inducible Auxotrophy in Escherichia coli and Salmonella enterica Serovar Typhimurium by Using an Insertion Element and a Hyperactive Transposase. Appl Env Microbiol. 2006;72: 4717–4725. doi:10.1128/AEM.00492-06

53. Guiney DGMD, Fierer JMD. The Role of the spv Genes in Salmonella Pathogenesis. Front Microbiol. 2011;2. doi:10.3389/fmicb.2011.00129

54. Lesnick ML, Reiner NE, Fierer J, Guiney DG. The Salmonella spvB virulence gene encodes an enzyme that ADP-ribosylates actin and destabilizes the cytoskeleton of eukaryotic cells. Mol Microbiol. 2001;39: 1464–1470. doi:10.1046/j.1365-2958.2001.02360.x

55. Lee E-J, Groisman EA. An antisense RNA that governs the expression kinetics of a multifunctional virulence gene. Mol Microbiol. 2010;76: 1020–1033. doi:10.1111/j.1365-2958.2010.07161.x

56. Alix E, Blanc-Potard A-B. MgtC: a key player in intramacrophage survival. Trends Microbiol. 2007;15: 252–256. doi:10.1016/j.tim.2007.03.007

57. Berg DE, Weiss A, Crossland L. Polarity of Tn5 insertion mutations in Escherichia coli. J Bacteriol. 1980;142: 439–446.

58. Wang A, Roth JR. Activation of Silent Genes by Transposons Tn5 and Tn10. Genetics. 1988;120: 875–885.

59. Pati NB, Vishwakarma V, Jaiswal S, Periaswamy B, Hardt W-D, Suar M. Deletion of invH gene in Salmonella enterica serovar Typhimurium limits the secretion of Sip effector proteins. Microbes Infect. 2013;15: 66–73. doi:10.1016/j.micinf.2012.10.014

60. Hancock SJ, Phan M-D, Peters KM, Forde BM, Chong TM, Yin W-F, et al. Identification of IncA/C Plasmid Replication and Maintenance Genes and Development of a Plasmid Multilocus Sequence Typing Scheme. Antimicrob Agents Chemother. 2017;61: e01740–16. doi:10.1128/AAC.01740-16

61. Phan MD, Forde BM, Peters KM, Sarkar S, Hancock S, Stanton-Cook M, et al. Molecular Characterization of a Multidrug Resistance IncF Plasmid from the Globally Disseminated Escherichia coli ST131 Clone. PLOS ONE. 2015;10: e0122369. doi:10.1371/journal.pone.0122369

62. Llanes C, Gabant P, Couturier M, Bayer L, Plesiat P. Molecular Analysis of the Replication Elements of the Broad-Host-Range RepA/C Replicon. Plasmid. 1996;36: 26–35. doi:10.1006/plas.1996.0028

63. Deighan P, Beloin C, Dorman CJ. Three-way interactions among the Sfh, StpA and H-NS nucleoid-structuring proteins of Shigella flexneri 2a strain 2457T. Mol Microbiol. 2003;48: 1401–1416. doi:10.1046/j.1365-2958.2003.03515.x

64. Beloin C, Deighan P, Doyle M, Dorman CJ. Shigella flexneri 2a strain 2457T expresses three members of the H-NS-like protein family: characterization of the Sfh protein. Mol Genet Genomics. 2003;270: 66–77. doi:10.1007/s00438-003-0897-0

65. Doyle M, Fookes M, Ivens A, Mangan MW, Wain J, Dorman CJ. An H-NS-like Stealth Protein Aids Horizontal DNA Transmission in Bacteria. Science. 2007;315: 251–252. doi:10.1126/science.1137550

66. Yamaichi Y, Chao MC, Sasabe J, Clark L, Davis BM, Yamamoto N, et al. High-resolution genetic analysis of the requirements for horizontal transmission of the ESBL plasmid from Escherichia coli O104:H4. Nucleic Acids Res. 2015;43: 348–360. doi:10.1093/nar/gku1262

67. Poidevin M, Sato M, Altinoglu I, Delaplace M, Sato C, Yamaichi Y. Mutation in ESBL Plasmid from Escherichia coli O104:H4 Leads Autoagglutination and Enhanced Plasmid Dissemination. Front Microbiol. 2018;9. doi:10.3389/fmicb.2018.00130

68. Dziewit L, Czarnecki J, Wibberg D, Radlinska M, Mrozek P, Szymczak M, et al. Architecture and functions of a multipartite genome of the methylotrophic bacterium Paracoccus aminophilus JCM 7686, containing primary and secondary chromids. BMC Genomics. 2014;15: 124. doi:10.1186/1471-2164-15-124

69. Ebert M, Laaß S, Burghartz M, Petersen J, Koßmehl S, Wöhlbrand L, et al. Transposon Mutagenesis Identified Chromosomal and Plasmid Genes Essential for Adaptation of the Marine Bacterium Dinoroseobacter shibae to Anaerobic Conditions. J Bacteriol. 2013;195: 4769–4777. doi:10.1128/JB.00860-13

70. Tazzyman SJ, Bonhoeffer S. Why There Are No Essential Genes on Plasmids. Mol Biol Evol. 2015;32: 3079–3088. doi:10.1093/molbev/msu293

71. Haseltine WA, Block R. Synthesis of Guanosine Tetra- and Pentaphosphate Requires the Presence of a Codon-Specific, Uncharged Transfer Ribonucleic Acid in the Acceptor Site of Ribosomes. Proc Natl Acad Sci U S A. 1973;70: 1564–1568.

72. Agirrezabala X, Fernández IS, Kelley AC, Cartón DG, Ramakrishnan V, Valle M. The ribosome triggers the stringent response by RelA via a highly distorted tRNA. EMBO Rep. 2013;14: 811–816. doi:10.1038/embor.2013.106

73. Chaliotis A, Vlastaridis P, Mossialos D, Ibba M, Becker HD, Stathopoulos C, et al. The complex evolutionary history of aminoacyl-tRNA synthetases. Nucleic Acids Res. 2017;45: 1059–1068. doi:10.1093/nar/gkw1182

74. Knuth K, Niesalla H, Hueck CJ, Fuchs TM. Large-scale identification of essential Salmonella genes by trapping lethal insertions. Mol Microbiol. 2004;51: 1729–1744. doi:10.1046/j.1365-2958.2003.03944.x

75. Kröger C, Colgan A, Srikumar S, Händler K, Sivasankaran SK, Hammarlöf DL, et al. An Infection-Relevant Transcriptomic Compendium for Salmonella enterica Serovar Typhimurium. Cell Host Microbe. 2013;14: 683–695. doi:10.1016/j.chom.2013.11.010

76. Martin M. Cutadapt removes adapter sequences from high-throughput sequencing reads. EMBnet.journal. 2011;17: 10–12. doi:10.14806/ej.17.1.200

77. Li H. Aligning sequence reads, clone sequences and assembly contigs with BWA-MEM. ArXiv13033997 Q-Bio. 2013; Available: http://arxiv.org/abs/1303.3997

78. Barquist L, Mayho M, Cummins C, Cain AK, Boinett CJ, Page AJ, et al. The TraDIS toolkit: sequencing and analysis for dense transposon mutant libraries. Bioinformatics. 2016;32: 1109–1111. doi:10.1093/bioinformatics/btw022

79. Vohra P, Chaudhuri RR, Mayho M, Vrettou C, Chintoan-Uta C, Thomson NR, et al. Retrospective application of transposon-directed insertion-site sequencing to investigate niche-specific virulence of Salmonella Typhimurium in cattle. BMC Genomics. 2019;20: 20. doi:10.1186/s12864-018-5319-0

80. Datsenko KA, Wanner BL. One-step inactivation of chromosomal genes in Escherichia coli K-12 using PCR products. Proc Natl Acad Sci U S A. 2000;97: 6640–6645.

81. Kintz E, Davies MR, Hammarlöf DL, Canals R, Hinton JCD, van der Woude MW. A BTP1 prophage gene present in invasive non-typhoidal Salmonella determines composition and length of the O-antigen of the lipopolysaccharide. Mol Microbiol. 2015;96: 263–275. doi:10.1111/mmi.12933

82. O’Leary NA, Wright MW, Brister JR, Ciufo S, Haddad D, McVeigh R, et al. Reference sequence (RefSeq) database at NCBI: current status, taxonomic expansion, and functional annotation. Nucleic Acids Res. 2016;44: D733–D745. doi:10.1093/nar/gkv1189

83. Ashton PM, Owen SV, Kaindama L, Rowe WPM, Lane CR, Larkin L, et al. Public health surveillance in the UK revolutionises our understanding of the invasive Salmonella Typhimurium epidemic in Africa. Genome Med. 2017;9. doi:10.1186/s13073-017-0480-7

